# Rab11-exosome biogenesis regulators mediate Aβ-induced intercellular propagation of endolysosomal trafficking defects and neurodegeneration

**DOI:** 10.64898/2026.09.17.752336

**Authors:** Amy Cording, Bhavna Verma, Adam Arous, Shauna Rice, Harrison Vine, Lewis Blincowe, S. Mark Wainwright, Preman J. Singh, Deborah C. I. Goberdhan, Clive Wilson

## Abstract

Alzheimer’s Disease (AD) is characterised by two histopathological hallmarks, intracellular tau-containing neurofibrillary tangles and extracellular amyloid plaques containing β-amyloid (Aβ), a specific cleavage product of Amyloid Precursor Protein (APP). Initiation of Aβ-induced neuronal pathology, however, has been postulated to involve intracellular events affecting endolysosomal trafficking that can propagate between cells. Recent studies in non-neuronal *Drosophila* secondary cells (SCs) have revealed that Aβ interferes with APP-regulated protein aggregation events, which package signalling molecules into insoluble dense-core granules (DCGs) during normal regulated secretion, inducing endolysosomal defects that are transferred to other cells. Here, using SCs, we show that knockdown of the gene encoding accessory ESCRT-III protein Chmp5, which selectively regulates the formation of intraluminal vesicles (ILVs) inside SC DCG compartments that are subsequently released as Rab11-exosomes, inhibits propagation of Aβ-induced endolysosomal trafficking defects. Furthermore, knocking down *Chmp5* or other *accessory ESCRT-III* genes also suppresses Aβ-induced, neurodegeneration-dependent morphological defects in the *Drosophila* eye. We conclude that genes controlling the Rab11-exosome biogenesis pathway play a key role in both Aβ-induced endolysosomal trafficking defects associated with aberrant regulated secretion and cellular events leading to neurodegeneration. Our findings indicate these processes are linked and may suggest new target pathways for future development of AD therapeutics.

## Introduction

Alzheimer’s Disease (AD) is the leading cause of dementia worldwide, predicted to affect more than 100 million people by 2050 (Gustavsson et al., 2023; Liu et al., 2026). Intracellular hyperphosphorylated tau-containing neurofibrillary tangles (Rawat et al., 2022) and extracellular amyloid plaques, composed of Aβ-peptides, specific cleavage products of Amyloid Precursor Protein (APP) (Weglinski and Jeans, 2023), are key histopathological AD hallmarks. However, there remains considerable controversy concerning the cell biological triggers that initiate degeneration.

One appealing model, the ‘traffic jam hypothesis’ (Kimura and Yanagisawa, 2018), postulates that defects in trafficking through the endosomal and lysosomal system lead to accumulation of intracellular compartments that contain Aβ, which fails to be degraded. This endolysosomal phenotype can be propagated to other cells and is an initiating step in disrupting synaptic trafficking, structure and integrity.

Hyperphosphorylated tau, which is associated with neurofibrillary tangles, appears to interact with Aβ and pathological Aβ-oligomers via endolysosome-mediated mechanisms (Miyoshi et al., 2021; Schützmann et al., 2021; Gao et al., 2025), suggesting one potential explanation for how tau and Aβ are functionally linked in disease. However, until recently, the question of whether AD-related endolysosomal defects might reflect disruption of a specific trafficking process involved in normal APP processing has remained unresolved.

We recently studied the functions of APP in *Drosophila* secondary cells (SCs) (Singh et al., 2025), prostate-like cells of the male accessory gland (MAG) that contain highly enlarged secretory and lysosomal compartments (Corrigan et al., 2014; Redhai et al., 2016; Fan et al., 2020). In these cells, dynamic events inside compartments, which accompany maturation and quality control steps in regulated secretion, have been visualised for the first time. In cells of higher organisms, proteins destined for secretion aggregate into dense-core granules (DCGs) inside regulated secretory compartments (Gondré-Lewis et al., 2012). These only dissipate when the compartments fuse with the plasma membrane or when defective compartments are targeted for lysosomal degradation in processes collectively termed crinophagy (Szenci et al., 2023). In SCs, the *Drosophila* orthologue of transmembrane APP, APPL, is required to promote normal DCG protein aggregation (Singh et al., 2025). It must then be cleaved, probably by β-secretase, which makes the first cleavage required for Aβ generation, to dissociate DCG aggregates from the compartment’s limiting membrane. APPL’s functions in regulated secretion can be substituted by human APP, suggesting an evolutionarily conserved role in this process (Singh et al., 2025).

When either wild type Aβ or amyloidogenic mutant forms of Aβ associated with early-onset familial AD are expressed in the secretory pathway of SCs, APP processing is disrupted, DCGs do not form normally and more compartments are targeted for lysosomal degradation (Singh et al., 2025). However, this targeting is aberrant and partially degraded compartments accumulate within SCs, generating an endolysosomal trafficking defect. Furthermore, the contents of these abnormal compartments appear to be secreted and build up inside endolysosomes of other neighbouring non-Aβ-expressing cells that endocytose them, propagating the endolysosomal phenotype (Singh et al., 2025). These findings suggest that Aβ-induced endolysosomal pathologies in AD can result from aberrant regulated secretion (Verma et al., 2026), a hypothesis supported by *in silico* analysis of human AD-associated genes (Kuo et al., 2021).

Maturation of regulated secretory compartments requires a transition of compartments emerging from the *trans*-Golgi network (TGN) to recycling endosomal identity, which can be tracked by the progressive acquisition of recycling endosomal marker, Rab11 (Wells et al., 2023; Stockhammer et al., 2024). Regulated secretory compartments in SCs generate intraluminal vesicles (ILVs), a process that also appears to be conserved in human DCG compartments (Wang et al., 2025), and these vesicles are secreted as specialised Rab11-exosomes (Fan et al., 2020). Previous studies have suggested that APPL is trafficked to these ILVs (Singh et al., 2025) and that knocking down genes encoding specific components of the four Endosomal Sorting Complex Required for Transport (ESCRT) complexes, which regulate ILV biogenesis, disrupts DCG formation in SCs (Marie et al., 2023). Furthermore, transfer of Aβ and its oligomers from neurons to other cells, including other neurons, appears to involve exosomes (Eitan et al., 2016; Tong et al., 2024). We therefore postulated that Rab11-exosomes might be involved in propagating Aβ-induced endolysosomal trafficking defects between cells.

Here we show that the endolysosomal defects induced by Aβ-expressing SCs spread throughout the epithelial cells of the MAG. Knockdown of *Chmp5*, which encodes an accessory ESCRT-III protein that selectively regulates Rab11-exosome formation (Marie et al., 2023), strongly suppresses this propagation. Knockdown of *Chmp5* and other *accessory ESCRT-III* genes also suppresses the neurodegeneration-associated morphological defects induced by Aβ expression in the fly eye. Our findings therefore suggest that the defects in endolysosomal trafficking caused by Aβ-induced disruption of regulated secretion may play important roles in AD-related degeneration and that these might be mitigated by inhibiting the Rab11-exosome biogenesis machinery.

## Results

### SC-expressed human Aβ induces endolysosomal defects in epithelial cells throughout the MAG

Previously, we showed that a GFP-tagged form of the DCG protein MFAS, the fly orthologue of mammalian extracellular matrix protein TGF-β-induced (TGFBI), when expressed from the endogenous gene locus, is selectively produced at high levels in SCs and is packaged into SC DCGs (Singh et al., 2025). When Aβ is expressed in the approximately 40 SCs at the distal tip of each lobe in the bi-lobed MAG, uptake of secreted GFP-MFAS into adjacent main cells (MCs) is enhanced and GFP accumulates in enlarged endolysosomal compartments, which stain with LysoTracker Red (Singh et al., 2025). This suggests that these compartments cannot degrade endocytosed DCG proteins normally and have a higher luminal pH, which fails to fully quench GFP fluorescence.

Our original analysis of Aβ-induced MC endolysosomal defects was restricted to cells directly adjacent to SCs. For these MCs, the endocytosed GFP-MFAS released by SCs could have either been secreted into the MAG lumen or transferred between cells across their lateral membranes. To confirm that luminal GFP-MFAS is responsible for the propagation phenotype, we investigated the uptake of GFP-MFAS by epithelial cells throughout the MAG. Male flies were generated in which adult SCs expressed either wild type Aβ (Aβ-wt) or the amyloidogenic Aβ-Iowa mutant (Grabowski et al., 2001), fused with a cleavable N-terminal ER signal sequence to target them to the secretory pathway (Chouhan et al., 2016; Metsla et al., 2022; Singh et al., 2025). Adult-only expression solely within the SCs of MAGs was achieved using *dsx-GAL4* activation of UAS-regulated transgenes in the presence of the temperature-sensitive GAL4 inhibitor, GAL80^ts^, shifting from 25°C to 29°C, which destabilises GAL80^ts^, at eclosion (Fan et al., 2020).

As previously reported (Singh et al., 2025), Aβ expression affected DCG biogenesis and endolysosomal trafficking in SCs, leading to increased accumulation of partially acidified compartments and lysosomes, and in the case of Aβ-Iowa, frequent incomplete aggregation of proteins into a large DCG (Figure S1A-C). Analysis of the MC epithelial layer in the central part of dissected *ex vivo* MAGs (Figure 1A) indicated that expressing Aβ isoforms in SCs increased the size of MC acidic compartments and particularly for Aβ-Iowa, led to accumulation of more GFP-MFAS inside these enlarged compartments (Figure S1D-F). This mirrored our previous findings for the distal part of the MAG and suggested endolysosomal defects propagate throughout the gland.

**Figure 1.**
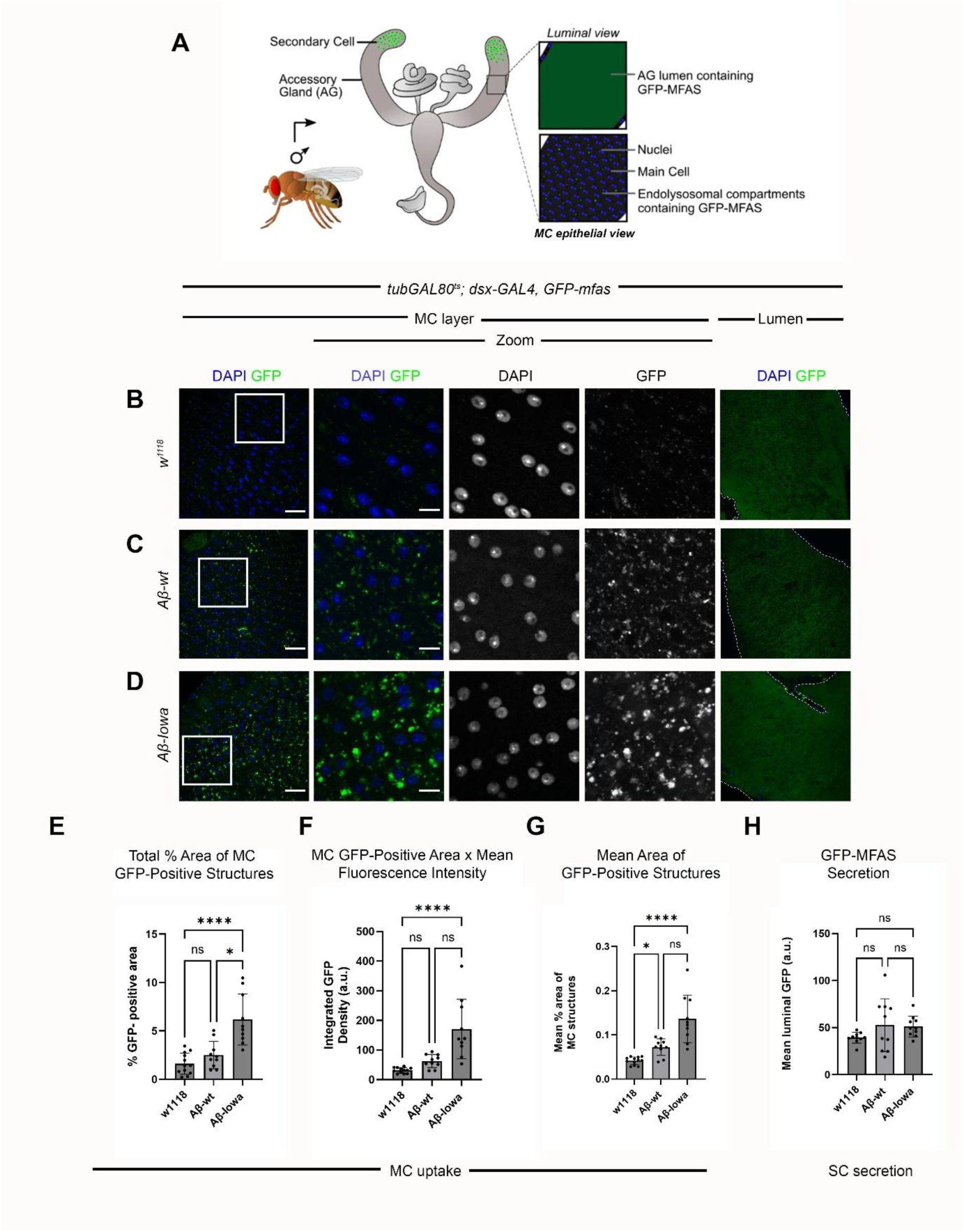
SC-expressed human Aβ induces endolysosomal defects in epithelial cells at the centre of the MAG. **A.** Schematic showing experimental strategy. SCs are only located at the distal end of the MAG, but secrete GFP-MFAS and other DCG compartment contents into the gland’s lumen, from where they can be endocytosed by MCs. **B-D.** Confocal images of transverse sections through the MC epithelial layer in the middle of the MAG and through the MAG lumen (panels on far right) in MAGs from 6-day-old males expressing the *GFP-mfas* gene trap, and either no other transgene (**B**), or UAS-regulated Aβ-wt (**C**) or Aβ-Iowa (**D**) specifically in SCs. Zoom panels show 3-fold magnification. **E-H.** Bar charts showing total % area of GFP-MFAS fluorescence in MC layer in central part of MAG (**E**), integrated intensity (area multiplied by mean GFP fluorescence intensity in MCs) (**F**), mean % area of individual GFP-MFAS-labelled MC compartments/puncta (**G**) and mean GFP-MFAS fluorescence intensity in lumen of MAG (**H**). Note that SC-specific Aβ-wt and Aβ-Iowa expression induce larger GFP-MFAS-containing structures inside MCs, with Aβ-Iowa also significantly increasing total GFP-MFAS-containing area and integrated intensity inside MCs. White squares mark positions of Zoom areas. Scale bars: 30 µm and 10 µm for higher magnification (Zoom) views.

To permit routine quantification of these changes, we fixed MAGs from GFP-MFAS-expressing males, stained them with DAPI to locate the thin squamous MC epithelial layer and imaged GFP distribution by confocal microscopy (Figure 1A). This revealed increased uptake of GFP-MFAS by MCs in central and proximal regions of MAGs from 6-day-old males, when distal SCs expressed Aβ-wt and Aβ-Iowa versus non-expressing controls (Figure 1B-D; Figure S2A-C).

The endocytosed GFP-MFAS located in the middle of the MAG was quantified by measuring the total % area containing GFP fluorescence in a cross-section of the epithelial MCs identified by their nuclear staining, the integrated density (% area x mean fluorescence signal intensity), and the mean % area for each intracellular GFP-MFAS-containing region. All three of these were significantly increased in comparison to controls when SCs expressed Aβ-Iowa (Figure 1E-G). For MAGs in which Aβ-wt was expressed in SCs, there was also a clear increase in the mean % area of each GFP-MFAS-containing region (Figure 1G); for many, but not all, glands, the total % area of fluorescence and the integrated intensity also appeared to be increased relative to the mean control values (Figure 1E and 1F), though because of greater variation, neither of these changes reached significance. For both Aβ genotypes, when compared to controls, the number of MC structures containing GFP and the mean fluorescent signal intensity were either unaffected, or in the case of Aβ-Iowa, slightly increased for fluorescence intensity (Figure S2D and S2E). Overall, these findings confirm that Aβ expression in SCs induces expansion of MC endolysosomal compartments, which accumulate the labelled SC DCG cargo GFP-MFAS.

The central region of the MAG lumen was also imaged in these different genetic backgrounds and secreted GFP-MFAS fluorescence levels measured using a protocol modified from Singh et al. (2025) (Figure 1A-D and 1H). For Aβ-wt and Aβ-Iowa, more variable levels of fluorescence were observed than for controls; some glands contained high levels of GFP-MFAS, as we had previously reported (Singh et al., 2025); however overall, there was no significant increase in secretion as measured by this assay (Figure 1H), eliminating elevated secretion as an explanation for the endolysosomal changes in MCs.

To confirm that SC secretion of GFP-MFAS is required for Aβ-induced transfer to MCs, we co-expressed Dad, a negative transcriptional regulator of BMP signalling, in SCs, either with Aβ-wt or the Aβ-Iowa mutant (Figure 2). Autocrine BMP signalling stimulates SC luminal secretion and this is suppressed by Dad (Corrigan et el., 2014; Redhai et al., 2016). No adult males expressing both Aβ-wt and Dad eclosed, presumably because leaky developmental expression under the control of the *dsx-GAL4*/*GAL80^ts^* system leads to synthetic lethality. However, expression of Dad in SCs expressing Aβ-Iowa reduced the levels of GFP-MFAS in the MAG lumen and drastically decreased the uptake of this fusion protein into MCs (Figure 2).

**Figure 2.**
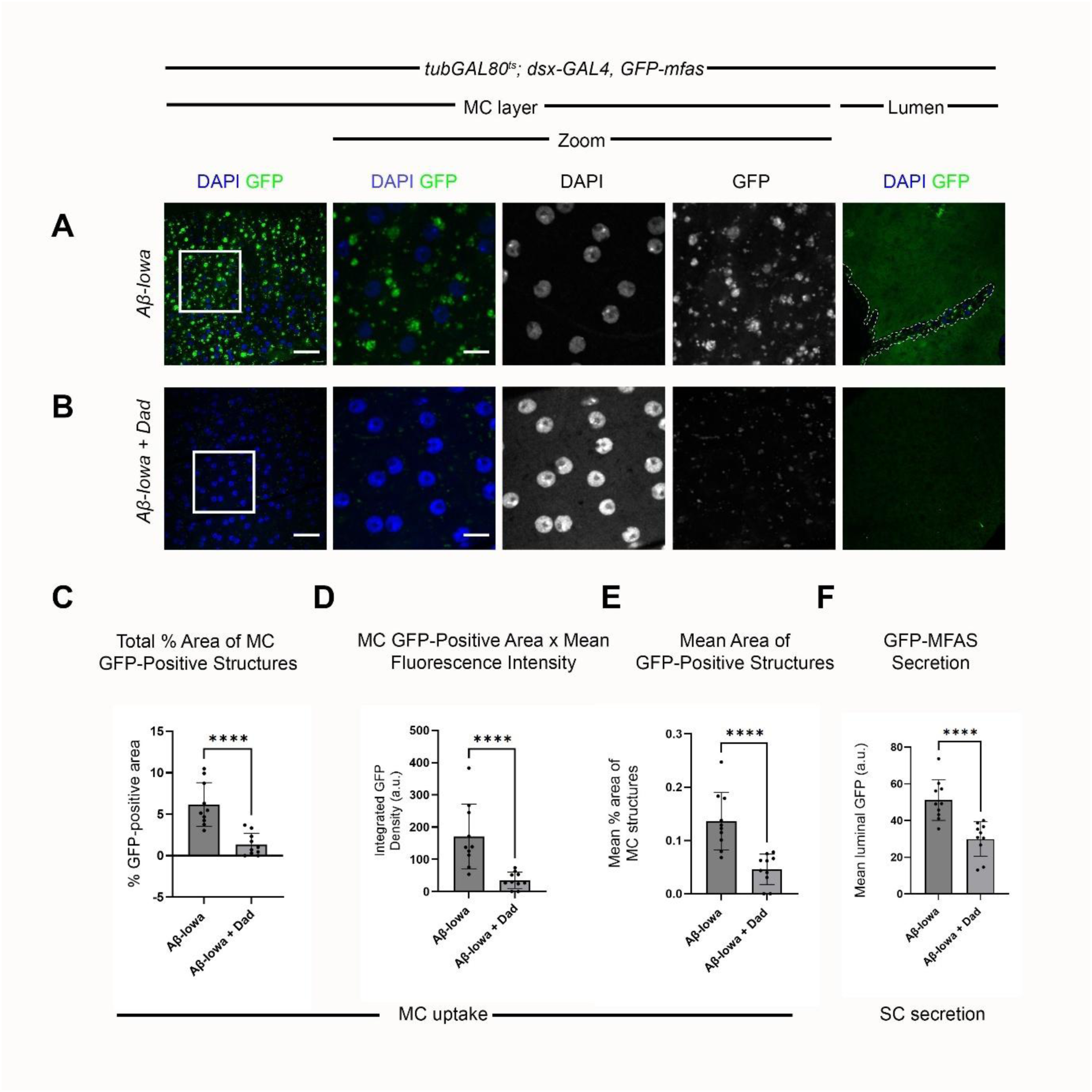
Aβ-Iowa-induced uptake of GFP-MFAS by MCs is dependent on secretion-inducing BMP signalling in SCs. **A, B.** Confocal images showing accumulation of GFP-MFAS in the MC layer and in the MAG lumen of glands in which SCs express Aβ-Iowa in the presence (**B**) or absence (**A**) of overexpressed Dad. Note the substantial reduction in secretion and MC GFP-MFAS when Dad is expressed. **C-F.** Bar charts showing total % area of GFP-MFAS fluorescence in MC layer in central part of MAG (**C**), integrated intensity of MC fluorescence (**D**), mean % area of individual GFP-MFAS-labelled MC compartments (**E**) and mean GFP-MFAS fluorescence intensity in lumen of MAG (**F**). White squares mark positions of Zoom areas. Scale bars: 30 µm and 10 µm for higher magnification (Zoom) views.

In summary, our data strongly support the idea that MCs endocytose DCG proteins secreted by SCs and that the uptake process is enhanced, and/or the degradation suppressed, when SCs express wild type or amyloidogenic mutant Aβ. This leads to enlarged MC endolysosomal compartments that contain increased levels of undegraded fluorescent GFP-MFAS that is not quenched, probably because the pH of these defective compartments is elevated.

### Aβ isoforms integrate into DCGs in maturing DCG compartments

The effects of Aβ expression in SCs on the transfer of GFP-MFAS and propagation of endolysosomal defects to MCs is consistent with our previous observations: Aβ affects the aggregation of GFP-MFAS and presumably other secreted DCG proteins in SCs (Singh et al., 2025), potentially via aberrant membrane:aggregate interactions, and this increases uptake of these proteins by MCs following secretion. To investigate whether Aβ-peptides directly incorporate into the DCG, we used a previously validated antibody for these peptides, 6E10 (Ray et al., 2017). This antibody cross-reacts with human Aβ-peptides and APP, but not *Drosophila* APPL. The antibody produced high level background staining around the muscular capsule of the MAG (Figure 3A), making it difficult to assess Aβ localisation in the thin MC layer. However, both Aβ-wt and Aβ-Iowa strongly co-localised with most GFP-MFAS-positive DCGs inside SCs, although the intensity of antibody staining varied considerably between different DCGs (Figure 3).

**Figure 3.**
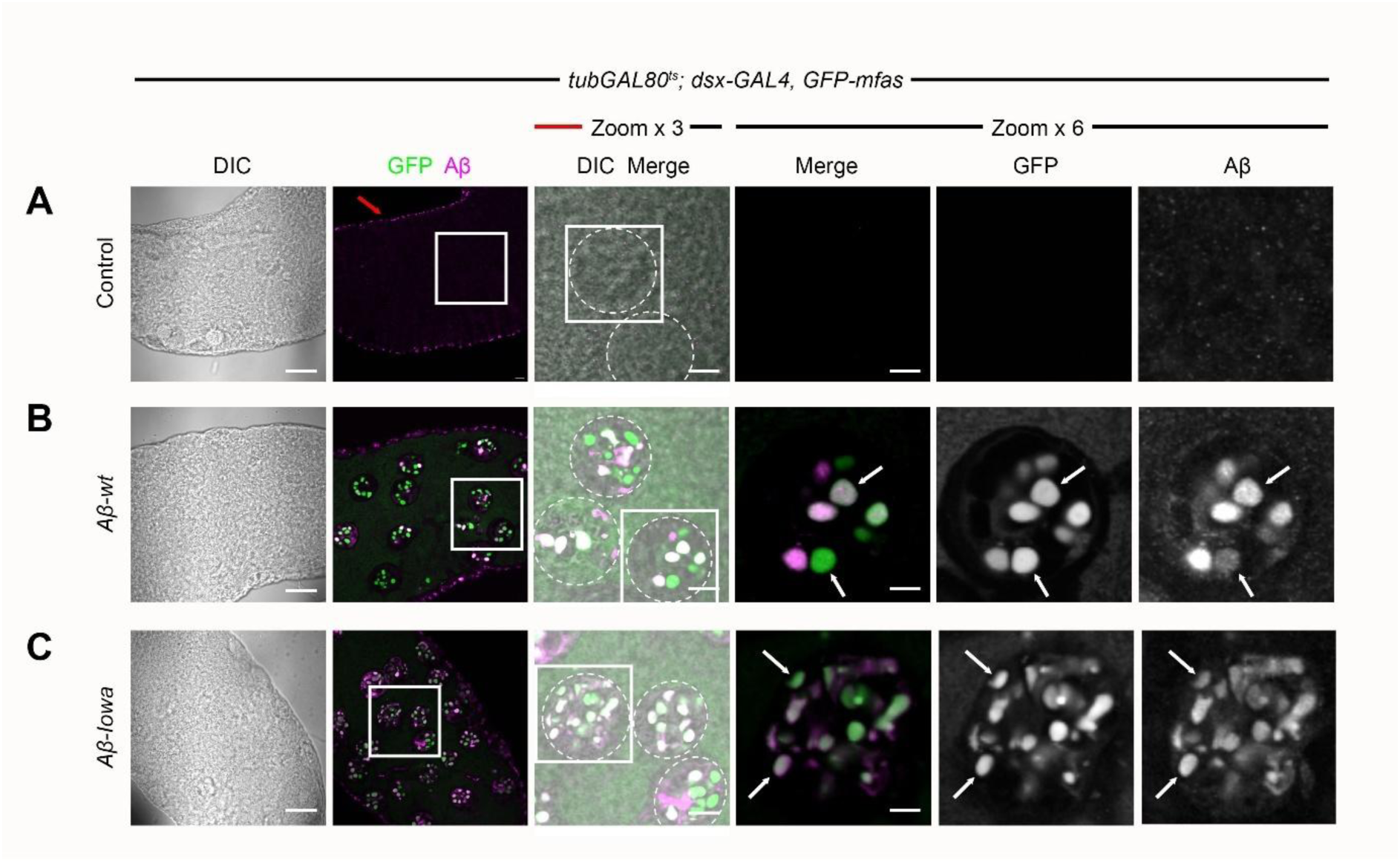
SC-expressed Aβ-wt and Aβ-Iowa localises to DCGs. **A-C.** Confocal images of distal parts of wild type MAGs (**A**) and MAGs carrying the *GFP-mfas* gene trap and expressing either Aβ-wt (**B**) or Aβ-Iowa (**C**) in SCs, which have all been immunostained with an antibody that cross-reacts with a human Aβ epitope. Note that generally Aβ staining is observed in almost all SC DCG compartments and overlaps with GFP-MFAS-labelled DCGs (white arrows), although the fluorescence level is more variable than for GFP. The anti-Aβ antibody also cross-reacts inappropriately with structures in the peripheral muscle layer, even in MAGs that do not express Aβ (red arrow). Example SCs are outlined with white dashed circle. White squares mark positions of Zoom areas. Scale bars: 30 µm and 10 µm and 5 µm for higher magnification views (Zoom x 3 and Zoom x 6 respectively).

We conclude that Aβ expressed in SCs is incorporated into DCGs and alters DCG compartment maturation. When secreted by these cells, Aβ’s effects on the DCG protein MFAS either persist or are re-established in the MAG lumen, so that MFAS is abnormally endocytosed by MCs, disrupting their endolysosomal trafficking.

### SC exosomes are endocytosed by MCs together with DCG proteins

SC DCG compartments also generate ILVs that are secreted into the MAG lumen as Rab11-exosomes (Fan et al., 2020; Marie et al., 2023). We investigated whether these exosomes are endocytosed by MCs, and whether this process is affected by SC Aβ expression. We co-expressed an mCherry-labelled form of the human exosome marker CD63, which marks Rab11-exosomes (Fan et al., 2020), in SCs expressing GFP-MFAS, so that we could easily identify GFP-MFAS-containing compartments, as well as endocytosed exosomes, in MCs. We could not generate males in which GFP-MFAS, and the CD63-mCherry and Aβ-Iowa transgenes, all of which reside on the same chromosome as *dsx-GAL4*, were co-expressed. However, CD63-mCherry localised inside MCs, both in control MAGs and in MAGs containing Aβ-wt-expressing SCs, frequently co-localising with GFP-MFAS (Figure 4). Furthermore, as for the GFP-MFAS signal (Figures 1B-D and S1D-F), Aβ-wt expression led to more diffuse distribution of CD63-mCherry in MCs, consistent with the enlargement of endolysosomal compartments in these cells. Therefore, endocytosis of SC-derived exosomes into MCs accompanies GFP-MFAS. It occurs in wild type MAGs, but also in MAGs in which MC endolysosomes have been disrupted by SC-specific Aβ expression.

**Figure 4.**
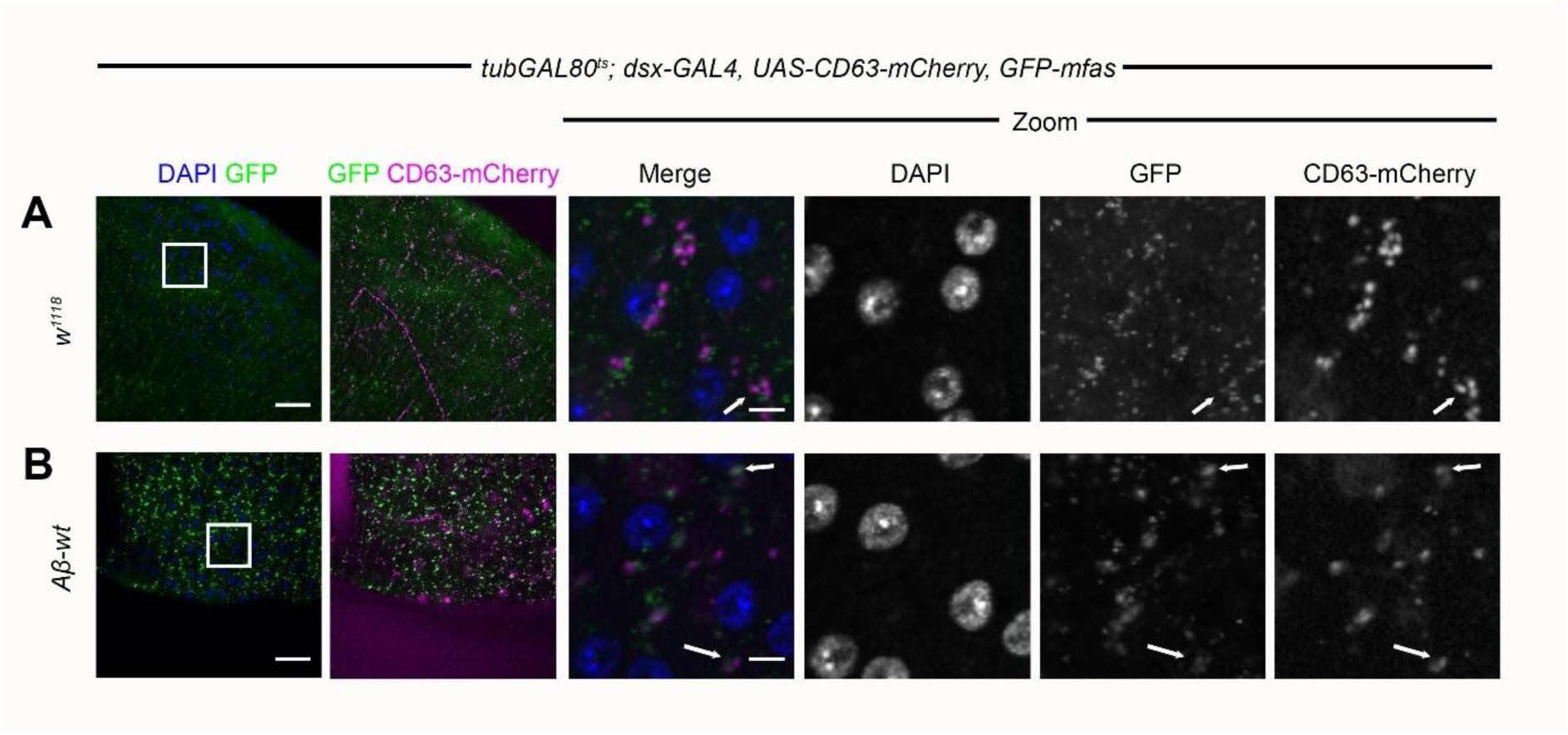
SC Rab11-exosomes are also endocytosed by MCs. **A, B.** Confocal images of central regions of MAGs from males carrying the *GFP-mfas* gene trap and expressing *CD63-mCherry* in SCs, either with no other transgene (**A**) or together with Aβ-wt (**B**). Note that CD63-mCherry is endocytosed by MCs under both conditions and much of it co-localises with GFP-MFAS (arrows) or is in close proximity. The GFP-MFAS signal is more diffuse when SCs express Aβ-wt (see also Figures 1 and S1D-F) and this is mirrored in the distribution of CD63-mCherry, consistent with localisation in larger endolysosomes. White squares mark positions of Zoom areas. Scale bars: 30 µm and 10 µm for higher magnification (Zoom) views.

### Inhibiting expression of Rab11-exosome biogenesis regulator, Chmp5, in SCs suppresses Aβ-induced propagation of endolysosomal defects to MCs

Our previous analysis suggested that Aβ expression in SCs suppresses the dissociation of aggregated DCG proteins from membranes, an important step in normal DCG maturation, and that the resulting defective DCG compartments fail to traffic normally to lysosomes and are aberrantly secreted (Singh et al., 2025). ILVs in DCG compartments are associated with DCGs both during and after the compartment maturation process (Fan et al., 2020; Dar et al., 2021; Wells et al., 2023; Singh et al., 2025). Indeed, these ILVs appear to play a role in DCG biogenesis (Marie et al., 2023; Singh et al., 2025). We hypothesised that Rab11-exosomes could promote Aβ-induced endolysosomal trafficking defects in MCs because of the stabilised interactions between membranes and GFP-MFAS-containing protein aggregates in Aβ-expressing SCs might persist outside these cells.

To test this, we inhibited Rab11-exosome biogenesis in SCs in the presence and absence of Aβ-peptides. ILV biogenesis in DCG compartments is controlled by the core components of the four ESCRT complexes, which can also generate late endosomal exosomes; however, we have previously shown in fly and human cells that accessory ESCRT-III proteins are selectively involved in Rab11-exosome formation versus exosome biogenesis in late endosomes (Marie et al., 2023). Knockdown of genes encoding these proteins suppresses ILV formation in DCG compartments, but does not detectably affect the ILV-dependent degradation of ubiquitinated proteins in late endosomes and lysosomes, unlike core ESCRT knockdowns (Marie et al., 2023).

We therefore assessed the transfer of GFP-MFAS to MCs following SC knockdown of the *accessory ESCRT-III* gene, *Chmp5*. When we previously expressed a control RNAi that targets the eye pigmentation gene *rosy*, which encodes the enzyme xanthine dehydrogenase, in SCs, we found that it increased GFP-MFAS levels in MCs (Singh et al., 2025). We therefore used an alternative RNAi targeting a non-*Drosophila* gene, *mCherry*; in the absence of Aβ, it also appeared to enhance MC GFP-MFAS uptake when compared to non-expressing controls (compare Figures 5A, 5C-E to Figure 1), but to a lesser extent than *rosy-RNAi*. This result and other studies suggest that activating the RNAi processing machinery in SCs may enhance the intercellular transfer of GFP-MFAS. Despite this enhancement, knockdown of *Chmp5* strongly suppressed levels of internalised MC GFP-MFAS, as measured by all three assays we employed, not only in MAGs where SCs expressed wild type Aβ or Aβ-Iowa, but even in non-Aβ-expressing cells (Figure 5). This manipulation also reduced the mean GFP fluorescence intensity of the MC endolysosomal compartments, and for MAGs in which SCs did not express Aβ or expressed Aβ-wt, the numbers of large endolysosomal structures detected in MCs were also decreased (Figure S3). These findings indicate that *Chmp5* knockdown in SCs strongly reduces GFP-MFAS endocytosis and/or enhances its degradation in MCs.

**Figure 5.**
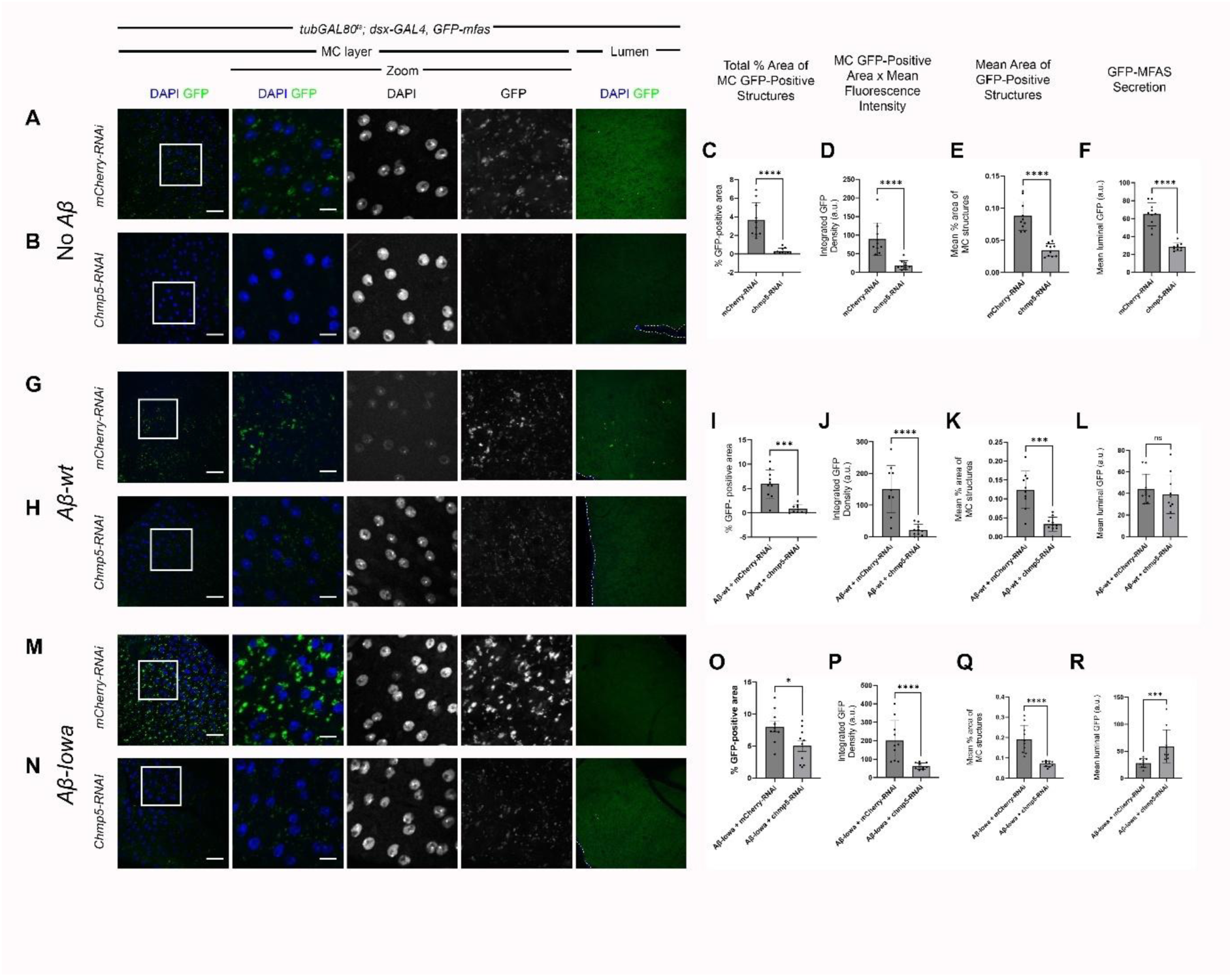
SC-specific *Chmp5* knockdown strongly suppresses Aβ-induced uptake of GFP-MFAS into MCs. **A**, **B**, **G**, **H**, **M**, **N.** Confocal images of central regions of MAGs from males carrying the *GFP-mfas* gene trap and expressing different knockdown constructs in SCs, as well as Aβ-wt (**G**, **H**) and Aβ-Iowa (**M**, **N**) constructs. In the absence of Aβ expression, SC-specific *Chmp5* knockdown reduces levels of GFP-MFAS in MCs (**B**), when compared to an *mCherry* knockdown control (**A**). GFP-MFAS levels in MCs are also reduced by *Chmp5* knockdown, when compared to *mCherry* knockdown controls, in MAGs where SCs express either Aβ-wt (**G**, **H**) or Aβ-Iowa (**M**, **N**). **C-F**, **I-L**, **O-R.** Bar charts quantifying *Chmp5* knockdown-induced suppression of GFP-MFAS transfer to MCs in genotypes where SCs express no Aβ (**C-F**), Aβ-wt (**I-L**) or Aβ-Iowa (**O-R**). Significant reductions are observed for all genotypes using assays measuring total % area of GFP-MFAS fluorescence in MC layer in central part of MAG (**C**, **I**, **O**), integrated density (area multiplied by mean GFP fluorescence intensity in MCs) (**D**, **J**, **P**), and mean % area of individual GFP-MFAS-labelled MC compartments (**E**, **K**, **Q**), following *Chmp5* knockdown. *Chmp5* knockdown also decreases GFP-MFAS secretion (mean GFP-MFAS fluorescence intensity in lumen of MAG) in MAGs where SCs do not express Aβ (**F**), but has no effect when Aβ-wt is expressed and appears to increase secretion in the presence of Aβ-Iowa. White squares mark positions of Zoom areas. Scale bars: 30 µm and 10 µm for higher magnification (Zoom) views.

Assessment of GFP-MFAS levels in the MAG lumen revealed that *Chmp5* knockdown did not reduce GFP-MFAS secretion by Aβ-wt-expressing SCs (Figures 5H and 5L) and indeed, seemed to increase it for Aβ-Iowa (Figures 5N, 5R), suggesting that altered MC uptake is not explained by decreased luminal GFP-MFAS. In control glands, *Chmp5* knockdown did induce a reduction in GFP-MFAS secretion (Figure 5F), though the inhibitory effect on MC uptake appeared stronger (Figures 5C-E). Overall, these data suggest that Rab11-exosomes and/or their ILV precursors play a critical role in the cell-to-cell transfer of DCG components and the propagation of Aβ-induced endolysosomal defects.

Some, but not all, *ESCRT* knockdowns inhibit the production of large DCGs in SCs, which might increase the targeting of these compartments to lysosomes (Marie et al., 2023) and therefore produce changes in the endolysosomal defects induced in Aβ-expressing cells. We therefore investigated whether in addition to its effects on phenotypic propagation, *Chmp5* knockdown altered endolysosomal trafficking in the SC quality control pathway for secretion or modified the endolysosomal trafficking defects observed in SCs expressing Aβ-wt or Aβ-Iowa (Figure 6).

**Figure 6.**
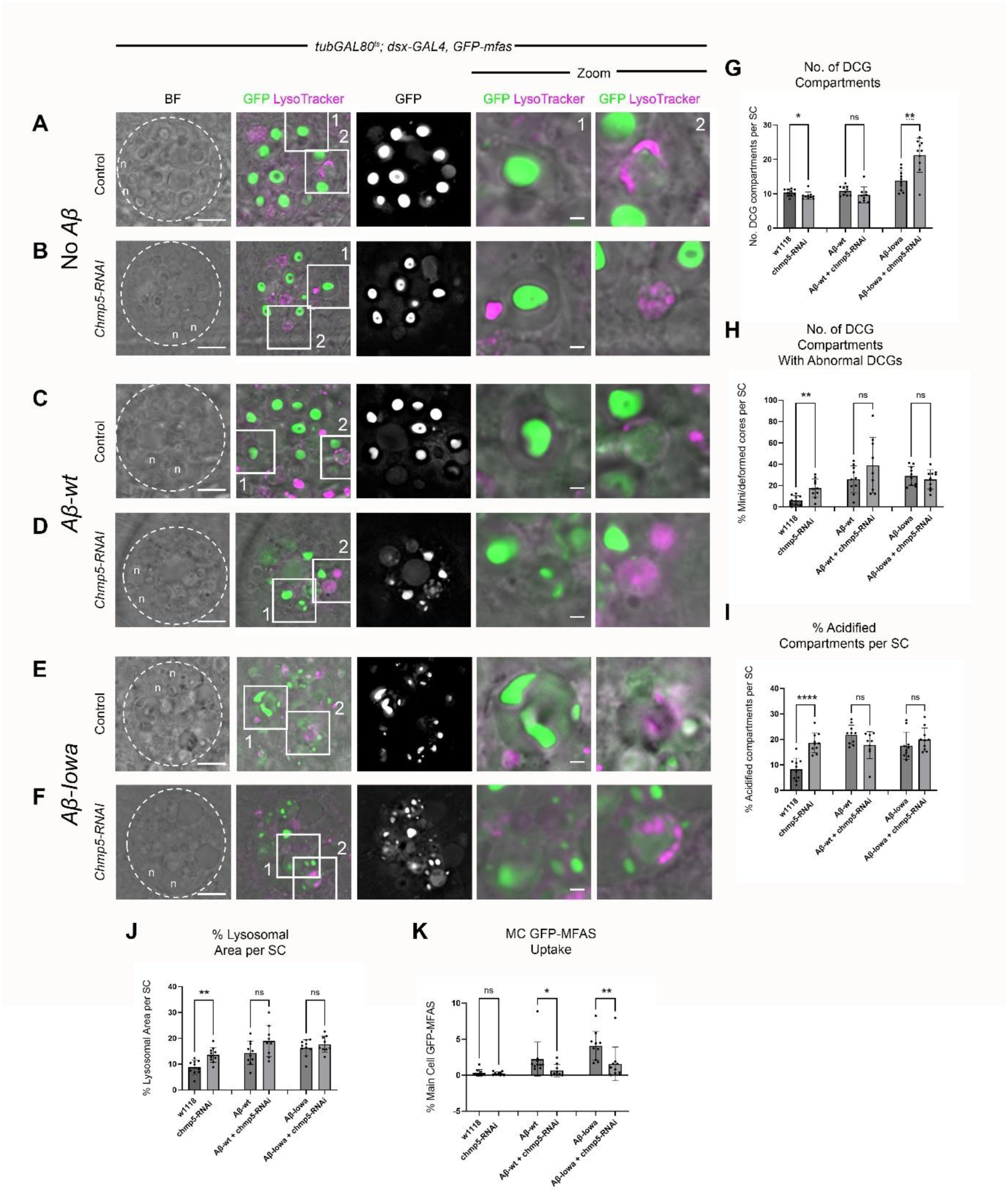
SC-specific *Chmp5* knockdown has only modest effects on DCG compartment biology and quality control. **A-F.** *Ex vivo* wide-field fluorescence images of single SC (marked by dashed circle). Adult SCs express no UAS-regulated transgene (**A**, **B**), Aβ (**C**, **D**) or Aβ-Iowa (**E**, **F**) in the presence (**B**, **D**, **F**) or absence (**A, C**, **E**) of a *UAS-Chmp5-RNAi* construct. These glands were dissected from males that also carry the *GFP-mfas* gene trap and were stained with LysoTracker Red. Bright-field view reveals outlines of large secretory and endolysosomal compartments, as well as the two nuclei (n) of these bi-nucleate cells. Zoom images show a DCG compartment (1) and an acidified DCG compartment (2). Apical views of SCs are shown; much flatter squamous MCs in deeper z-sections are not visible. **G-K.** Bar charts comparing SC DCG compartment number (**G**), % compartments with mini-cores or other deformed DCGs (**H**), % of acidified compartments (**I**), % lysosomal (LysoTracker Red-positive) area (**J**), and % MC area containing GFP-MFAS (**K**). White squares mark positions of Zoom areas. Scale bars: 10 µm and 2 µm for higher magnification (Zoom) views.

When expressed in SCs in the absence of Aβ, *Chmp5*-RNAi increased the number of acidified compartments and the lysosomal area in these cells (Figure 6A, 6B, 6I, 6J), perhaps because the cells contained a higher proportion of abnormal DCGs (Figure 6H). There was also a trend towards reduced uptake of GFP-MFAS by MCs compared to controls (Figure 6K), although this assay was insufficiently sensitive to detect a significant change, unlike the uptake assay in the middle of the MAG (Figure 5F).

When co-expressed with Aβ-wt or Aβ-Iowa (Figure 6C-F), a reduction in MC GFP-MFAS uptake was observed (Figure 6K, cf. Figures 5H-K and 5N-Q). However, there was no obvious reduction in abnormal DCGs, acidified compartments or lysosomal area, although DCG compartment number was increased in Aβ-Iowa-expressing cells (Figure 6G-J). Therefore, no cell-autonomous phenotypic suppression of Aβ-induced DCG defects by *Chmp5* knockdown in SCs could be detected by our standard assays.

Essentially all compartments in SCs that contain DCGs are Rab11-positive (Redhai et al., 2016; Fan et al. 2020), and this is not affected either by *Chmp5* knockdown (Marie et al., 2023) or Aβ isoform overexpression (Singh et al., 2025). Using flies carrying an N-terminal YFP protein fusion in the *Rab11* gene locus, we confirmed that although there was some minor fluctuation in DCG compartment number (Fig. S4G), for all the genotypes screened, their DCG compartments were marked with Rab11 (Figure S4A-F). This indicates that the step in the DCG maturation process involving transition to Rab11-positive recycling endosomal identity occurs normally (Figure S4).

YFP-Rab11 also labels a subset of ILVs inside these compartments (Fan et al., 2020). We have previously scored the proportion of Rab11-positive compartments containing fluorescent puncta as a measure of ILV biogenesis (Marie et al., 2023). This assay confirmed that *Chmp5* knockdown reduced the proportion of compartments containing ILVs when compared to controls in the presence and absence of Aβ-wt (Figures S4A-D and S4H). For Aβ-Iowa-expressing SCs, it was difficult to count all Rab11-positive compartments containing ILVs, because most of these ILVs are located next to the limiting membrane and appear as concentrations of YFP (Figure S4E and S4H). *Chmp5* knockdown increased the number of more centrally located ILVs (Figure S4I), explaining why it slightly increased the number of ILV-containing DCG compartments in this genetic background (Figure S4F and S4H).

Therefore, knocking down *Chmp5* in SCs selectively reduces the formation of ILVs in DCG compartments, but any subsequent effects on maturation and quality control of these compartments when Aβ isoforms are overexpressed were not detectable using the standard assays that we employ. However, this knockdown strongly suppresses the transfer of GFP-MFAS to other epithelial cells and the propagation of Aβ-induced endolysosomal defects to these cells. Interestingly, because *Chmp5* knockdown appears to inhibit transfer even from non-Aβ-expressing SCs, this exosome-dependent process may function physiologically, but it is strongly enhanced in the presence of Aβ-isoforms.

### Knockdown of *accessory ESCRT-III* genes suppresses Aβ-induced neurodegeneration in the eye

Our studies indicate that expressing wild type or mutant Aβ in SCs induces at least three phenotypes that might be relevant to initiating events in AD: disruption of aggregated protein dissociation from membranes in DCG compartments, endolysosomal trafficking defects, and propagation of these defects to other cells through SC secretion and endocytosis by MCs. *Chmp5* knockdown, which inhibits the formation of ILVs in DCG compartments and therefore the secretion of Rab11-exosomes (Marie et al., 2023), strongly suppresses the propagation phenotype, but only has more subtle, if any, effects on the Aβ-induced SC DCG and endolysosomal defects.

We have postulated that defects in regulated secretion induced by Aβ might initiate the neurodegenerative phenotypes observed in AD (Singh et al., 2025; Verma et al., 2026). Therefore, genetic manipulations that suppress any of these defects in SCs might also inhibit degeneration triggered by expression of Aβ in neurons.

To test this hypothesis, we employed a neurodegeneration model in the fly eye, which is commonly used to study AD-associated neurodegeneration. Expression of either wild type Aβ or Aβ-Iowa in the developing *Drosophila* eye using the eye-specific GMR-GAL4 driver disrupts the normal hexagonal array of ommatidial unit eyes, due to cell degeneration during the time that adult eye morphology is established (Tare 2011; Prüßing et al., 2013; Burnouf et al., 2015; Na et al., 2023). Several genetic and pharmacological suppressors of AD-associated neurodegeneration, as well as modifiers of secretion, partially rescue these phenotypes (Cao et al., 2008; Wang et al., 2015; De Mena et al., 2020; Deshpande et al., 2023; Na et al., 2023). Furthermore, Rab11 is known to play a critical role in apical trafficking and secretion in *Drosophila* photoreceptors, consistent with the idea that these cells, like many other cell types, have the basic regulated secretory biogenesis machinery present in SCs.

We confirmed that expression of Aβ-wt and Aβ-Iowa under the control of two different GMR-GAL4 insertions produced a disorganised eye phenotype (Figure 7). Previous analytical tools have been developed to study rough eye phenotypes (Diez-Hermano et al., 2015; Iyer et al., 2016; Tran et al., 2025), though we found that these did not work well with our stereomicroscopic images. We therefore adapted features of these methods and developed an automated ommatidial analysis pipeline to measure the regularity of the eye’s hexagonal array for approximately 250 central ommatidia. This determined the angle to the six closest ommatidia for each unit eye in control and Aβ-expressing eyes within a central elliptical region within the eye. We established a reference curve for each GMR-GAL4 line when crossed to flies carrying no UAS-transgene insertion; it contained six clear peaks, each separated by a 60 degree angle (Figure S6). When compared to this reference curve, we found that the irregular arrangement of individual Aβ-expressing eyes could be clearly distinguished from eyes of flies expressing a *lacZ* control transgene (Figures S5I).

**Figure 7.**
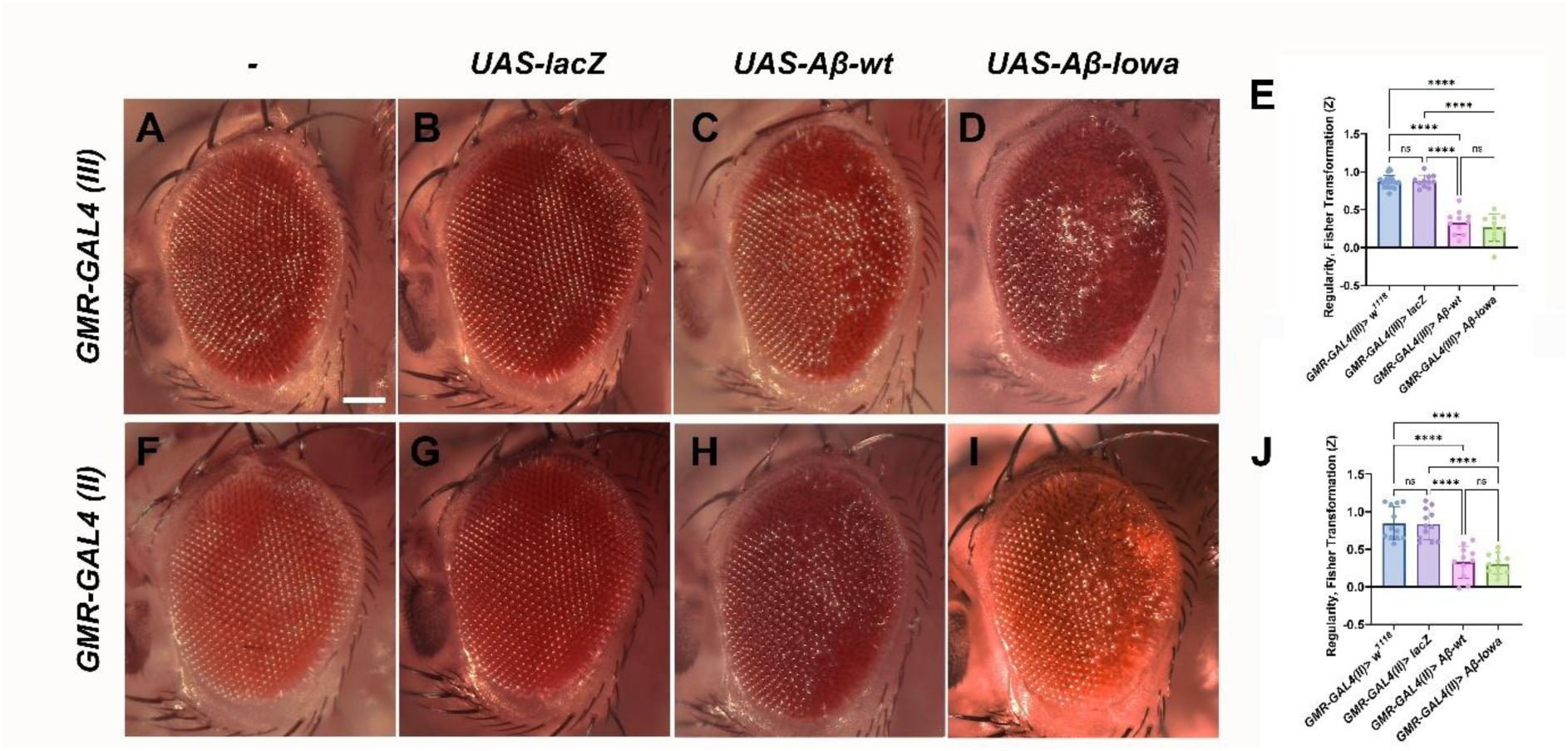
Expression of human Aβ-wt and Aβ-Iowa induce quantifiable disruption of ommatidial regularity in the *Drosophila* eye. **A-D.** Stereomicroscopic images of eyes from 2-4-day-old females carrying the third chromosome *GMR-GAL4 (III)* insertion in the presence of no other transgene (**A**), *UAS-lacZ* (**B**), *UAS-Aβ-wt* (**C**) and *UAS-Aβ-Iowa* (**D**). **E.** Bar chart plotting regularity score (as a Fisher Z transformation) for eyes of these genotypes relative to control *GMR-GAL4 (III)* reference curve. **F-I.** Stereomicroscopic images of eyes from 2-4-day-old females carrying the second chromosome *GMR-GAL4 (II)* insertion in the presence of no other transgene (**F**), *UAS-lacZ* (**G**), *UAS-Aβ-wt* (**H**) and *UAS-Aβ-Iowa* (**I**). **J.** Bar chart plotting regularity score (as a Fisher Z transformation) for eyes of these genotypes relative to control *GMR-GAL4 (II)* reference curve. Scale bar: 100 µm, applies to all images.

Expression of a control RNAi, *mCherry*-RNAi, had no effect either on the regularity of ommatidia in control eyes or on the irregularity induced by wild type Aβ and Aβ-Iowa (Figures S6, 8A, 8B, 8D, 8E, 8F and 8H). By contrast, *Chmp5*-RNAi suppressed Aβ-induced disorganisation in eyes expressing wild type Aβ, but not in Aβ-Iowa-expressing eyes (Figures 8C, 8D, 8G and 8H). When expressed alone, it induced some ommatidial irregularity with the third chromosomal *GMR-GAL4 (III)* driver (Figures S6A, S6B, S6F, S6G, S6H and S6L), suggesting that accessory ESCRT-III function might play physiological roles in regulating normal eye morphology during development.

**Figure 8.**
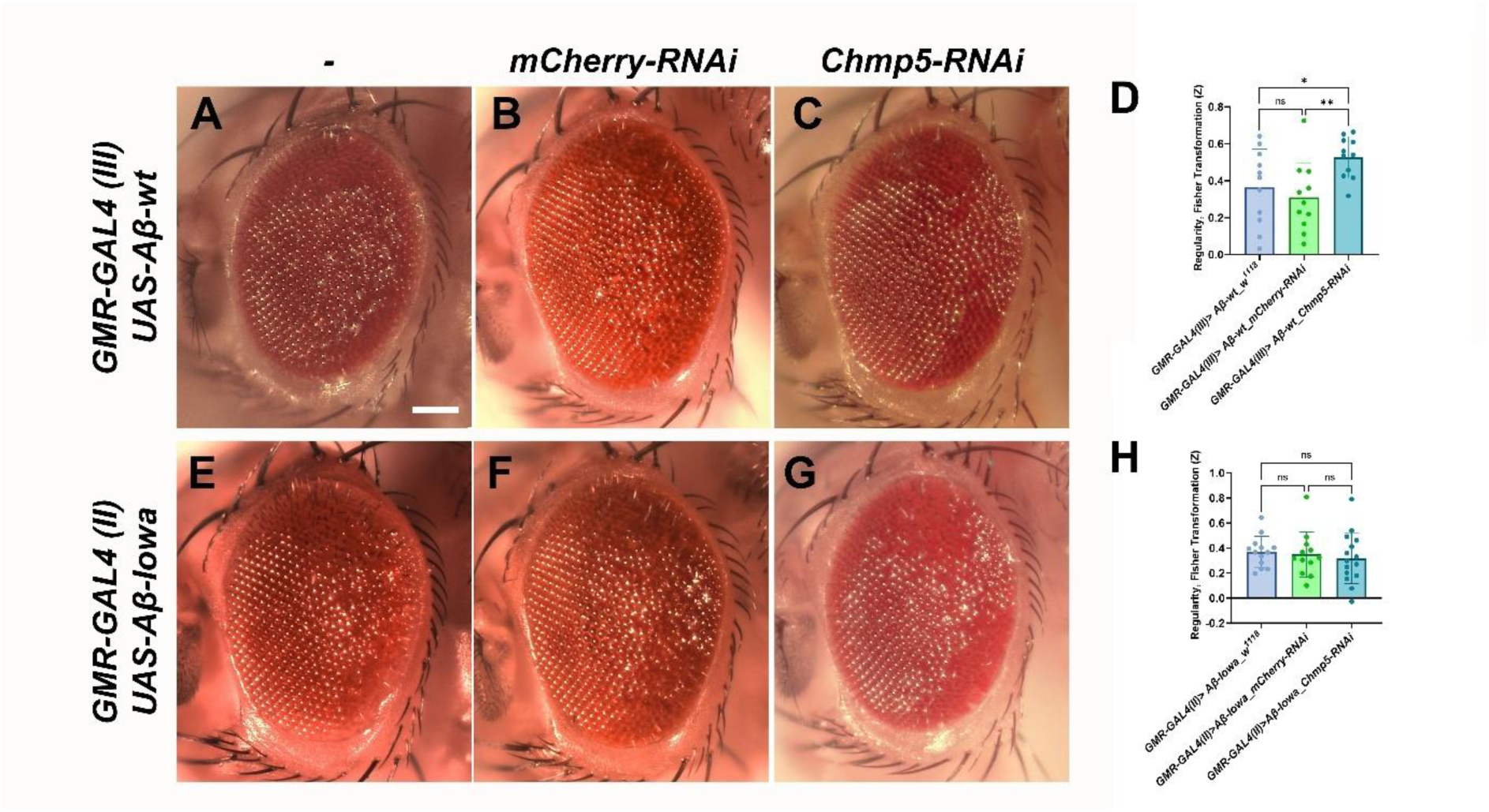
*Chmp5* knockdown suppresses Aβ-wt-induced disorganisation in the eye. **A-C.** Stereomicroscopic images of eyes from 2-4-day-old females carrying the third chromosome *GMR-GAL4 (III)* insertion in the presence of *UAS-Aβ-wt* and co-expressing either no other transgene (**A**), *UAS-mCherry-RNAi* (**B**) or *UAS-Chmp5-RNAi* (**C**). **D.** Bar chart plotting regularity score (as a Fisher Z transformation) for eyes of these genotypes relative to control *GMR-GAL4 (III)* reference curve. Note partial rescue of disorganisation by *Chmp5* knockdown relative to both controls. **E-G.** Stereomicroscopic images of eyes from 2-4-day-old females carrying the second chromosome *GMR-GAL4 (II)* insertion in the presence of *UAS-Aβ-Iowa* and co-expressing either no other transgene (**E**), *UAS-mCherry-RNAi* (**F**) or *UAS-Chmp5-RNAi* (**G**). **H.** Bar chart plotting regularity score (as a Fisher Z transformation) for eyes of these genotypes relative to control *GMR-GAL4 (II)* reference curve. No rescue is observed. Scale bar: 100 µm.

A previous study (Zhuang et al., 2023) has reported that RNAi-mediated depletion of the core ESCRTs *Stam*, *Vps25* and *Vps4* suppresses a degeneration phenotype induced by overexpressing human APP under GMR-GAL4 control in the fly eye, supporting the proposal that ESCRT function is required for degeneration in APP/Aβ-associated AD degeneration models. To confirm that the Rab11-exosome accessory ESCRT-III-mediated biogenesis pathway is an important player in Aβ-induced degeneration, we tested the effects of knocking down two other *accessory ESCRT-III* genes, *Ist1* and *Chmp1*, in the GMR-GAL4 model. Both knockdowns have previously been shown to selectively inhibit ILV formation in SC DCG compartments (Marie et al., 2023). They suppressed degeneration induced by both Aβ-wt and Aβ-Iowa (Figure 9), but had no effect when expressed alone in the eye (Figure S6).

**Figure 9.**
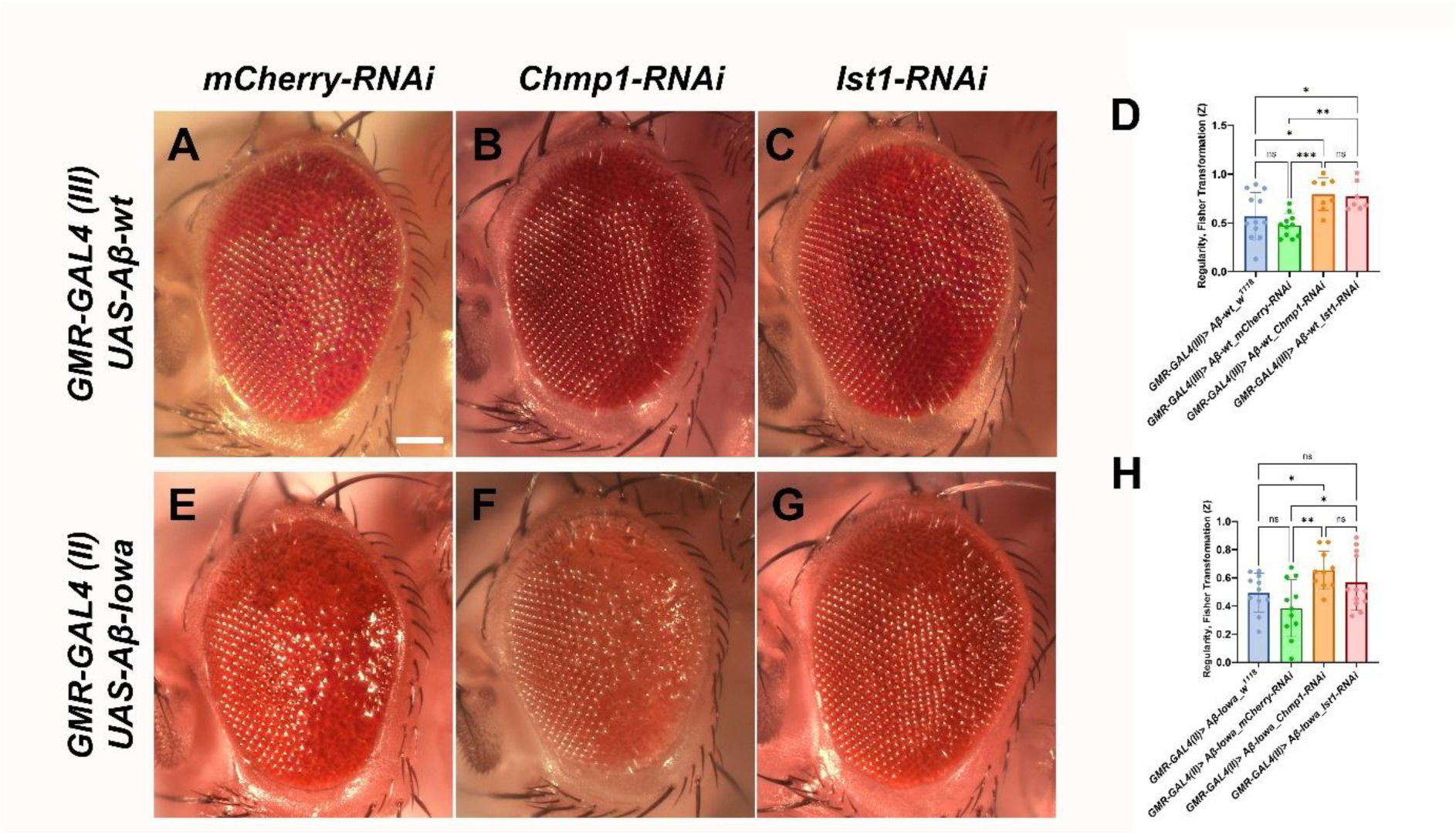
Knockdown of other *accessory ESCRT-III* genes suppresses Aβ-wt- and Aβ-Iowa-induced disorganisation in the eye. **A-C.** Stereomicroscopic images of eyes from 2-4-day-old females carrying the third chromosome *GMR-GAL4 (III)* insertion in the presence of *UAS-Aβ-wt* and co-expressing either *UAS-mCherry-RNAi* (**A**), or *UAS-Chmp1-RNAi* (**B**), or *UAS-Ist1-RNAi* (**C**). **D.** Bar chart plotting regularity score (as a Fisher Z transformation) for eyes of these genotypes relative to control *GMR-GAL4 (III)* reference curve. Note partial rescue of disorganisation by both *Chmp1* and *Ist1* knockdown relative to control. **E-G.** Stereomicroscopic images of eyes from 2-4-day-old females carrying the second chromosome *GMR-GAL4 (II)* insertion in the presence of *UAS-Aβ-Iowa* and co-expressing either *UAS-mCherry-RNAi* (**E**), or *UAS-Chmp1-RNAi* (**F**), or *UAS-Ist1-RNAi*. **H.** Bar chart plotting regularity score (as a Fisher Z transformation) for eyes of these genotypes relative to control *GMR-GAL4 (II)* reference curve. Note partial rescue of disorganisation by *Chmp1* and *Ist1* knockdown relative to RNAi-expressing control. Scale bar: 100 µm.

In summary, our data show that blocking the accessory ESCRT-III pathway, which is known to be selectively involved in Rab11-exosome biogenesis, suppresses the degeneration-associated morphological phenotype induced by different forms of Aβ. This suggests that the degenerative mechanism includes a process that involves Rab11-exosome biogenesis and potentially its downstream effects on the regulated secretory pathway that disrupt normal endocytic uptake of secreted DCG components.

## Discussion

The cell biological events by which the generation of Aβ from APP initiates neurodegeneration and promotes transfer of degenerative phenotypes to other cells still remain unclear. However, defects in endolysosomal trafficking and subsequent intercellular propagation of this phenotype appear to play important roles (Kimura and Yanagisawa, 2018). Our recent studies using *Drosophila* prostate-like SCs suggest that these defects may emerge from dysregulation of evolutionarily conserved APP-dependent processes controlling regulated secretion (Singh et al., 2025). These processes can generate Aβ normally, but at levels that are adequately handled by the cell’s quality control systems (Brothers et al., 2018; Verma et al., 2026).

Here, we show that expression in SCs of Aβ-wt and Aβ-Iowa, a mutant form of Aβ associated with early-onset familial AD (Grabowski et al., 2001), induces propagation of endolysosomal defects to epithelial cells throughout the *Drosophila* MAG. The aberrant endolysosomes produced contain excess levels of the DCG component MFAS as well as exosomes that are probably made in SC DCG compartments, though we cannot exclude that some CD63-mCherry-labelled vesicles may have been generated via other mechanisms. Genetically manipulating SCs to inhibit Rab11-exosome biogenesis has limited, if any, detectable effects on the DCG biogenesis pathway in Aβ-expressing cells, but strongly suppresses propagation. Furthermore, downregulating the Rab11-exosome biogenesis pathway also suppresses Aβ-induced degenerative effects in the fly eye, suggesting that disruption of processes linked to regulated secretion are involved in neurodegeneration and might be targets for future therapeutic strategies.

### Propagation of Aβ-induced endolysosomal trafficking defects are linked to the regulated secretory pathway

Taking advantage of the highly enlarged DCG compartments in SCs, we previously showed that Aβ expression disrupts the regulated secretory pathway and its quality control, and leads to endolysosomal defects that spread to neighbouring cells (Singh et al., 2025). For that study, it was possible to show using LysoTracker Red staining that the endolysosomal compartments in MCs are highly enlarged and that they exhibit enhanced GFP fluorescence derived from the GFP-tagged SC DCG protein MFAS. The MAG contains about 40 SCs at the distal end of each of its two lobes, which also contain about 1000 MCs. Here, we find that there is GFP accumulation inside enlarged MC endolysosomal compartments throughout the remainder of the MAG; for Aβ-Iowa, it was possible to show that as expected, the abnormal uptake is dependent on BMP signalling in SCs, which controls regulated secretion in these cells.

Immunostaining with a human Aβ-specific antibody demonstrated that Aβ-peptides are incorporated into SC DCGs. It was not possible to confirm whether these peptides are also transferred to MCs, because of the background staining produced by this antibody. However, our findings strongly support the idea that Aβ directly interferes with DCG biogenesis and compartment maturation, and raise the possibility that interactions with DCG components persist following secretion, explaining the aberrant uptake of secreted MFAS by MCs.

### Rab11-exosomes promote the intercellular propagation of Aβ-induced endolysosomal defects

Previous studies in rodent and human cells have revealed that Aβ and Aβ-oligomers are secreted in association with exosomes and that amyloid plaques contain exosomes and/or other extracellular vesicles (EVs), which promote their formation (Rajendran et al., 2006; Dinkins et al., 2016; Mowry et al., 2023; Gilbert et al., 2024). Furthermore, Aβ-associated EVs from AD patient plasma and cerebrospinal fluid (CSF), neural cells of AD patients, and plasma from rodent AD models, transfer pathological phenotypes to other neurons (Eitan et al., 2016). In addition, neurotoxic Aβ-oligomers have been reported to transfer between human neurons and glia via a mechanism involving exosomes (Tong et al., 2024). This process is stimulated by endolysosomal defects in the EV-secreting cells.

We reasoned that ILVs inside DCG compartments could represent at least some of the Aβ-associated EVs involved in propagation of endolysosomal defects. Indeed, *Chmp5* knockdown in SCs dramatically suppressed the abnormal uptake of GFP-MFAS by MCs, supporting this idea. We have previously shown that *accessory ESCRT-III* genes have a selective role in Rab11-exosome biogenesis in fly and human cells (Marie et al., 2023). Although we cannot exclude that the strong suppression of Aβ-induced propagation is associated with other cellular effects of *Chmp5* knockdown, this seems unlikely, since in this genetic background, GFP-MFAS is still secreted and no major changes are seen in the SC DCG and endolysosomal compartments mediating this secretion.

Several reports have highlighted important functions for Rab11-positive compartments in normal and pathological neuronal intercellular communication. For example, in flies, Rab11 is involved in pre-synaptic secretion at the neuromuscular junction, which includes release of exosome-like vesicles (Koles et al., 2012; Walsh et al., 2021). Studies in mammalian neurons also highlight a central role for Rab11 in the secretion of Aβ-associated exosomes in AD and AD models (Arbo et al., 2020). Indeed, there is an overlap between generic human Rab11-exosome signatures and proteins elevated in the cerebrospinal fluid of AD patients (Li et al., 2023; Singh et al., 2025).

Our findings suggest that aberrant intercellular transfer of DCG components is likely to be linked to the propagation of Aβ-induced exosome-associated pathologies. The enhanced transfer of these components may just reflect the increased endocytic activity or reduced degradative functions of MCs induced by Aβ. However, the observation that MFAS, membranes and ILVs interact abnormally in DCG compartments of Aβ-expressing SCs (Singh et al., 2025), raises the possibility that exosome complexes, coupled to DCG proteins as well as Aβ within the exosome’s ‘biomolecular corona’ (Paolini et al., 2026), could persist when secreted and elicit the endolysosomal defects in MCs.

Indeed, it remains unclear whether such a mechanism might be operating physiologically in the absence of elevated Aβ. Although *Chmp5* knockdown suppresses intercellular transfer of GFP-MFAS in SCs, which are not overexpressing Aβ, other RNAi-overexpressing controls exhibit increased intercellular GFP-MFAS transfer compared to non-expressing controls. It is therefore possible that hyperactivating the RNAi machinery is increasing intercellular propagation activity and *Chmp5* knockdown suppresses this increase. Nevertheless, Aβ has been implicated in physiological intercellular signalling (Brothers et al., 2018), so it is conceivable that this involves Rab11-exosome:aggregate complexes, an idea that should now be testable. SC-specific *Chmp5* knockdown inhibits a specific female response to mating, the rejection of other males (Marie et al., 2023), and it will be interesting to investigate whether *mfas*, a key regulator of DCG protein aggregation, APP or other DCG components are also involved in this exosome-mediated process.

### Knockdown of Rab11-exosome regulators also suppresses Aβ-induced neurodegeneration

GMR-GAL4-driven Aβ expression is well established to produce a morphological defect associated with neurodegeneration and has provided a powerful assay to identify suppressors of this process (Tare 2011; Prüßing et al., 2013; Burnouf et al., 2015; Na et al., 2023; Cao et al., 2008; Wang et al., 2015; De Mena et al., 2020; Deshpande et al., 2023; Na et al., 2023).

A study by Zhuang et al., 2023, reported that *core ESCRT* knockdown suppresses eye degeneration induced by GMR-GAL4-driven human APP. They argued that this manipulation reduced APP trafficking to late endosomes, where Aβ is normally generated. Our finding that *accessory ESCRT-III* knockdown, which we have shown in SCs and human colorectal cancer cells has a relatively selective effect on Rab11-exosomes versus late endosomal exosomes (Marie et al., 2023), can inhibit degeneration induced by pre-cleaved wild type and mutant Aβ suggests that other processes are involved in ESCRT-mediated suppression of neurodegeneration. One straightforward explanation emerging from our analysis in MAGs is that blocking propagation of endolysosomal defects by inhibiting Rab11-exosome biogenesis alleviates some of the effects of Aβ (see model in Figure 10). However, the GMR-GAL4 driver expresses Aβ in all photoreceptors, so phenotypic propagation from defective to normal photoreceptors would seem unlikely to play a major role in doing this.

**Figure 10.**
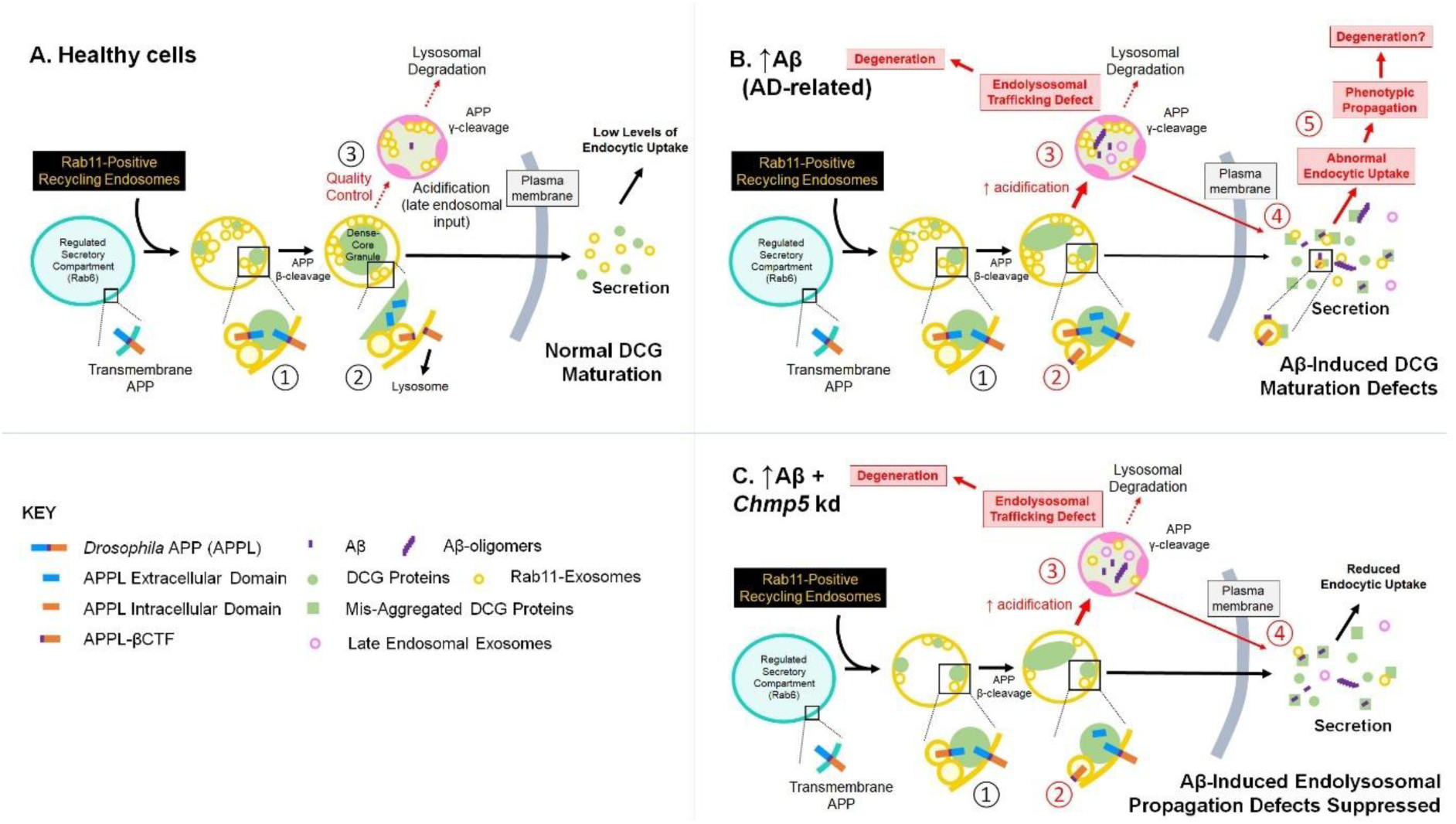
Proposed model to explain the role of Rab11-exosomes in intercellular propagation of Aβ-induced endolysosomal trafficking defects. Schematic shows the regulated secretory process in healthy cells (**A**), and in Aβ-expressing cells, which either produce normal (**B**) or reduced (**C**) levels of Rab11-exosomes. Regulated secretion involves DCG biogenesis (① and ②), a process requiring a compartmental switch to Rab11-positive recycling endosomal identity and APP processing, followed by quality control (③), and secretion. In healthy cells (**A**), only a small proportion of DCG compartments are targeted for lysosomal degradation, leading to minimal accumulation of Aβ. If Aβ starts to accumulate (**B**), this disrupts DCG biogenesis, causing reduced dissociation of membrane:aggregate interactions (eg. via APP; red ②) and increased, but aberrant, targeting of these compartments to lysosomes (red ③). Many of these compartments still appear to be secreted (red ④). The defective secreted material is abnormally endocytosed by other cells (as well as the secreting cells themselves [not shown]) and accumulates in enlarged endolysosomes, propagating the trafficking defect (red ⑤). This and the endolysosomal trafficking defects in secreting cells ultimately induce degeneration, both in SCs (Singh et al., 2025) and, as extensively reported (Kimura and Yanagisawa, 2018), in neurons. We find that *Chmp5* knockdown in SCs (**C**), which selectively inhibits Rab11-exosome biogenesis in human and *Drosophila* cells (Marie et al., 2023), does not have a detectable effect on Aβ-expressing cells. However, it suppresses the phenotypic propagation of endolysosomal defects, presumably by affecting the overall composition of aggregates and/or their interactions with vesicles, in secreting cells, or post-secretion (as shown here), or when the secreted material is endocytosed by other cells. This genetic manipulation is also sufficient to reduce Aβ-induced neurodegeneration in the fly eye, suggesting a role for Rab11-exosomes in Aβ-associated AD pathology.

An alternative possibility is that the uptake of material secreted from aberrantly assembled DCG compartments exacerbates endolysosomal trafficking defects in other Aβ-expressing cells, and potentially in the secreting cell itself. Although we did not observe any major changes in DCG or endolysosomal compartments of Aβ-expressing SCs in which *Chmp5* had been knocked down, these cells, like photoreceptors in the GMR-GAL4 assay, continue to synthesise more Aβ, even in the absence of endocytosis. They may, therefore, be less affected by the inhibition of the phenotypic propagation mechanisms than cells that require these mechanisms to initiate aberrant Aβ synthesis. This may also explain the incomplete suppression of neurodegeneration observed.

### Aberrant regulated secretion and an integrated model for initiation of AD pathology

If defective regulated secretion does play an important role in initiating Aβ-induced AD pathology and its propagation, where would other key players in AD, particularly tau, fit in? Aβ-associated endolysosomal defects have, in fact, been shown to induce tau pathologies in rodent neurons (Gao et al., 2025; Schützmann et al., 2021), while Aβ is implicated in the exosomal transfer of pathological tau seeds between human neurons (Miyoshi et al., 2021), processes that might be linked to aberrant secretory trafficking.

The activity-regulated cytoskeleton-associated protein, ARC, has recently also been directly implicated in tau seeding (Tyagi et al., 2026). Multiple studies link Arc to AD (Wu et al., 2011; Landgren et al., 2012; Bi et al., 2018) and in *Drosophila*, knockdown of the Arc orthologue, *Arc1*, suppresses tau-induced pathology in several AD models (Schulz et al., 2023). Since Arc is proposed to be intimately involved in the production of ILVs at pre-synaptic termini (Ashley et al., 2018) and can bind to the cytoplasmic domain of APP (Lee et al., 2023), it may provide one important link between the tau-modulated cytoskeleton and regulated secretory compartments that is relevant to the endolysosomal propagation phenotype. In this regard, it is interesting that *Arc1* is transcribed at particularly high levels in SCs and may therefore also play a role in this system (Immarigeon et al., 2021).

Our study raises the interesting possibility that the earliest pathological defects in AD are associated with secretion of abnormal Rab11-exosomes that carry disease-specific surface cargos (Figure 10). These exosomes are known to contain different protein payloads to late endosomal exosomes (Marie et al., 2023; Singh et al., 2025), so developing tests to distinguish neuronal Rab11-exosomes (called Rab11a-exosomes in humans) from other EVs in biofluids could provide a novel route to developing early-stage or even prodromal diagnostics tests. Recent analysis of the neuronal subcellular proteome in AD patients with dementia versus individuals with disease resilience suggests a co-ordinated reorganisation of endolysosomal proteins, particularly those associated with early and recycling endosomes, which is more robust in patients with cognitive decline (Jolly et al., 2026). It is tempting to speculate that Rab11a-exosomes may provide a read-out of these differences that will be relevant to prognosis.

The hypothesis that defects in regulated secretion and its quality control might lead to accumulation of endolysosomal compartments that generate Aβ, promoting protein aggregation and propagation of endolysosomal phenotypes between cells now needs to be tested in more detail in human and mammalian neurons (Verma et al., 2026). However, further studies in SCs should help to highlight additional players that disrupt DCG aggregation and promote the spreading of defectively aggregated material between cells, as well as genetic suppressors of these phenotypes. Importantly, if pathology in sporadic AD patients involves minor defects in DCG biogenesis that progressively accumulate and propagate through life, relatively subtle modulation of these suppressors might provide a therapeutic route at early stages that prevents the progression to dementia in later life.

## Materials and Methods

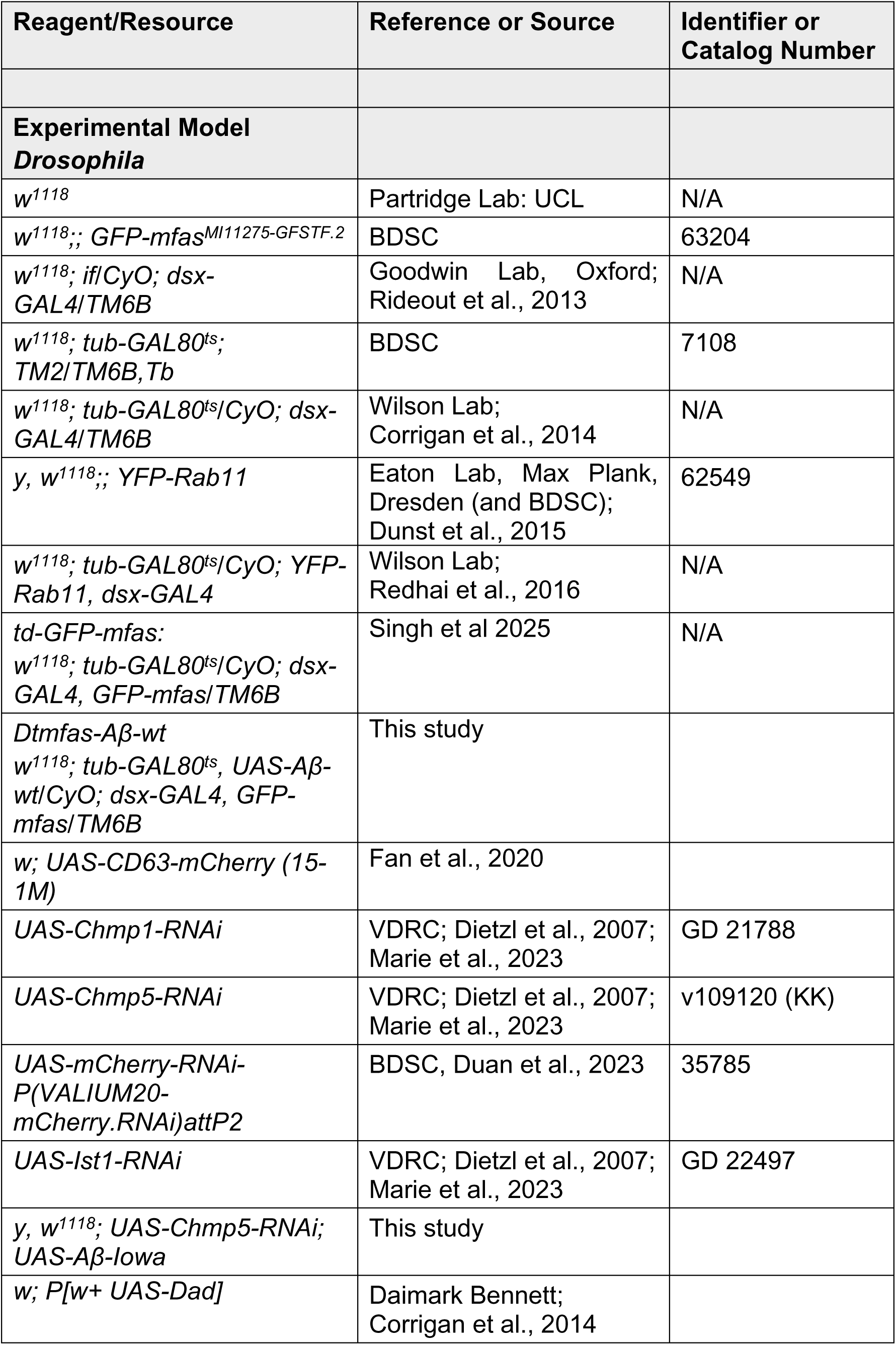
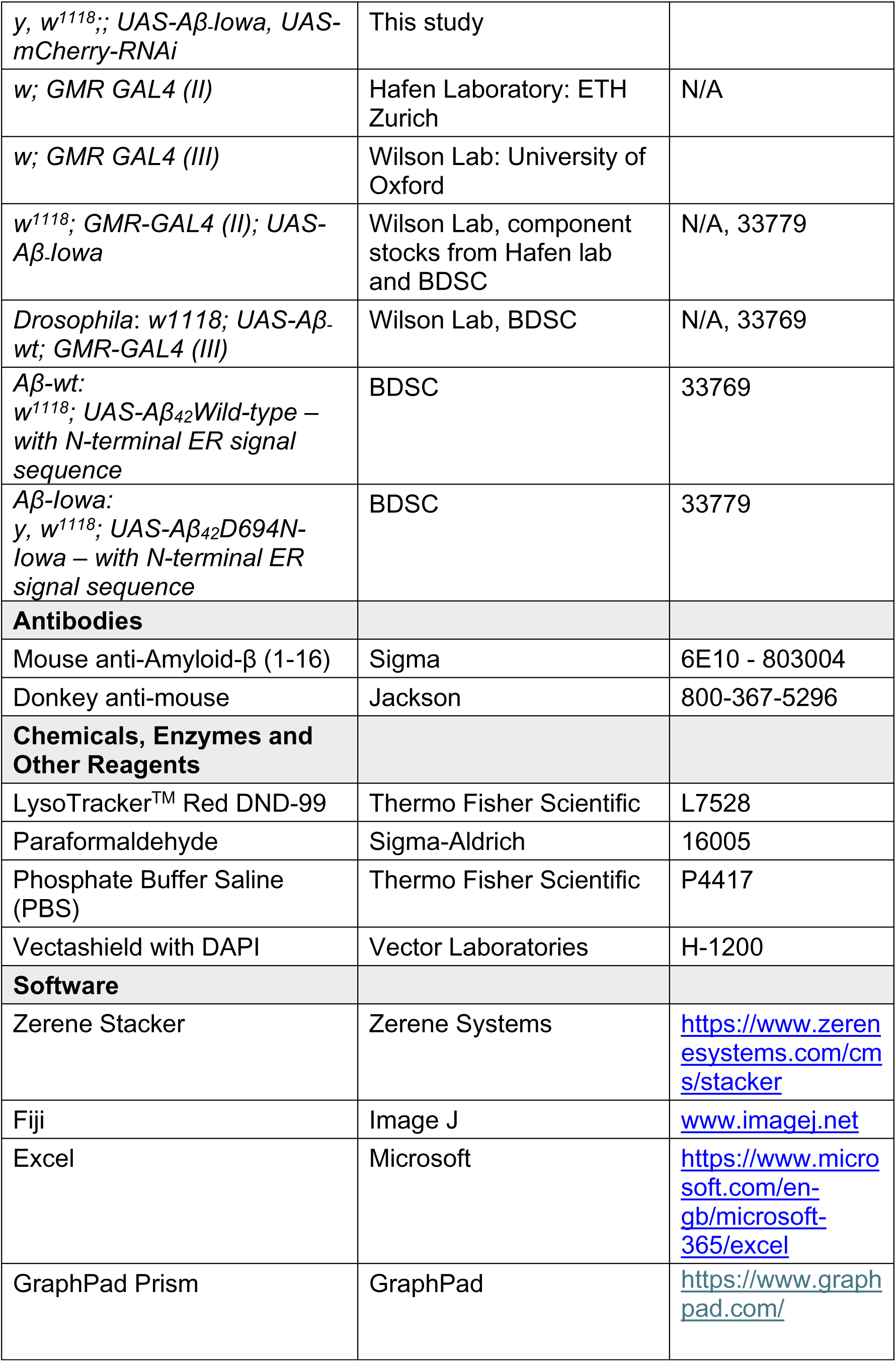

### Fly stocks and husbandry

*Drosophila* strains used in this manuscript are listed in the Reagents table. Most of the transgenic fly lines were obtained from Bloomington *Drosophila* Stock Centre (BDSC) and Vienna *Drosophila* Resource Centre (VDRC). Exceptions were as follows; *dsx-GAL4* (provided by S. Goodwin, Oxford, UK) (Rideout et al, 2010), YFP-*Rab11* fusion gene at endogenous *Rab* locus (provided by S. Eaton, Max Plank, Germany) (Dunst et al, 2015), UAS-Dad (provided by D. Bennett, University of Liverpool). In cases where lines were generated and/or used in previous studies, this is noted next to the stocks in the list below: *w^1118^* (provided by L.Partridge, UCL,UK) (eg. Singh et al., 2025): *GFP-mfas* gene trap (MI11275 GFSTF.2; BDSC 63204) (Nagarkar-Jaiswal et al, 2015; Singh et al., 2025): *UAS-CD63-mCherry* (Fan et al., 2020): *w; GAL80^ts^; dsx-GAL4* (Fan et al, 2020): *w; tub-GAL80^ts^; dsx-GAL4, GFP-mfas* (Singh et al., 2025): *w; tub-GAL80^ts^; YFP-Rab11, dsx-GAL4* (Fan et al, 2020): *UAS-Aβ-wt* (BDSC 33769) (Wu et al, 2017) and *UAS-Aβ-Iowa* (BDSC 33779; Vitruvean) (Chouhan et al, 2016) constructs contain a cleavable ER signal sequence, to direct them to the secretory system (used in Singh et al., 2025): *UAS-Aβ-wt* was combined with other transgenes to create *w; UAS-Aβ-wt, tub-GAL80^ts^; dsx-GAL4* and *w; UAS-Aβ-wt, tub-GAL80^ts^; dsx-GAL4, GFP-mfas*: *UAS-Chmp5-RNAi* (P{KK109120}VIE-260B; VDRC 101422; discontinued stock) (Dietzl et al., 2007; Marie et al, 2023): *UAS-mCherry-RNAi* (VALIUM20-mCherry; BDSC 35785) (Duan et al., 2023): *UAS-Chmp1-RNAi* (P{GD11219}v21788; VDRC 21788) (Dietzl et al., 2007; Marie et al., 2023): *UAS-Ist1-RNAi* ([P{GD11948}](https://flybase.org/reports/FBti0081469.html):VDRC 22497) (Dietzl et al., 2007; Marie et al., 2023): *w; GMR-GAL4 (II)* (provided by Hafen Laboratory: ETH Zurich) (Freeman, 1996), *GMR-GAL4 (III)* (generated in Wilson lab): *GMR-GAL4 (III*) was combined to create *w; UAS-Aβ-wt; GMR GAL4 (III)* and *GMR-GAL4 (II)* used to generate *w; GMR GAL4 (II); UAS-Aβ-Iowa*.

Flies were maintained at 25°C under a 12-hour light/dark cycle on standard cornmeal agar medium [12.5 g agar (F.Gutlind & Co. Ltd), 75 g cornmeal (B. T. P. Drewitt), 93 g glucose (Sigma-Aldrich, #G7021), 31.5 g inactivated yeast (Fermipan Red, Lallemand Baking), 8.6 g potassium sodium tartrate tetrahydrate (Sigma Aldrich, #S2377), 0.7 g calcium chloride dihydrate (Sigma-Aldrich, #21907), and 2.5g nipagin (Sigma-Aldrich, #H5501) dissolved in 12 ml ethanol, per litre]. They were transferred onto fresh food every 3-4 days.

For the experiments involving the MAG, female flies carrying the driver line *tub-GAL80^ts^; dsx-GAL4* alone or in combination with *GFP-mfas*, or *YFP-Rab11*, or *UAS CD63-mCherry* (in some cases, together with UAS-regulated forms of Aβ-wt or Aβ-Iowa) were crossed with male flies carrying UAS-transgenes or *w^1118^* (control), permitting a temperature-controlled induction of SC-specific target gene expression.

As in previous studies (Singh et al., 2025), for the experiments involving the MAG and live tissue imaging, virgin male offspring were collected upon eclosion and transferred to 29°C for 6 days to activate post-developmental SC-specific transgene expression.

For the experiments in which we quantified GFP-MFAS in the main cells, which involved imaging of fixed tissue, virgin males were transferred 24 h before eclosion to 29°C to allow time for RNAi to accumulate before DCG compartments begin to form in SCs, and then maintained at 29°C for 6 days post-eclosion.

For the experiments involving *Drosophila* eye analysis, female flies carrying *UAS-RNAi* transgenes or *w^1118^* controls were crossed with male flies carrying *GMR-GAL4* alone or in combination with *UAS-Aβ-wt* or *UAS-Aβ-Iowa*, to induce eye-specific expression of target genes. These crosses were maintained at 25°C throughout the whole experiment. Upon eclosion, female flies were separated from males and aged 2-4 days at 25°C before freezing and imaging.

### Preparation of MAGs for imaging

We used an adaptation of the methodology described by Fan et al., 2020 for live-cell imaging. Six-day-old adult male virgin flies were anaesthetised using CO_2_, then submerged in ice-cold 1X PBS (Thermo Fisher Scientific). The male reproductive system was carefully pulled out of the body cavity through the last abdominal segment with micro-forceps. The testes, seminal vesicles and ejaculatory bulb were left intact and attached to the gland, but fat tissues and the gut were gently removed to avoid tissue folding and interference during MAG imaging. The glands were then incubated with 500 nM Lysotracker Red (Thermo Fisher Scientific) in PBS for 2-5min on ice, followed by a wash with ice-cold PBS. The tissue was then stably mounted between two coverslips (rectangular: coverslip No.1, 22 mm×50mm, Fisher, #1237-3128, and round: coverslip No.1, 13 mm, #49492, VWR) in a drop of 1x PBS, with the set-up held together by a custom-built metal holder. Excess PBS was removed using a filter paper when the glands were ready for imaging, to slightly flatten the samples.

For confocal analysis, micro-dissections were performed in PBS at room temperature and then samples were fixed in 4% paraformaldehyde in PBS (Sigma-Aldrich) for 20 min. The glands were then washed in 1x PBS for approximately 5 min prior to mounting in a drop of Vectashield with DAPI (Vector Laboratories) on SuperFrost microscope slides (VWR). Samples were held in place with a coverslip (22 mm × 22 mm, 0.13–0.17 mm; Fisher)

### Antibody staining of fixed tissue

For antibody staining, MAGs were dissected and fixed in 4% paraformaldehyde (as described above). Fixed glands were then permeabilized for 6 × 10 min in PBST (1× PBS, 0.3% Triton X-100 [Sigma-Aldrich]), blocked for 30 min in PBSTG (PBST, 10% goat serum [Sigma-Aldrich]), and incubated overnight at 4°C with mouse anti-Aβ 6E10 (BioLegend, 39320) (Ray et al., 2017) diluted 1:500 in PBSTG. Glands were then washed for 6 × 10 min in PBST before incubation in a 1:400 dilution of Cy3– conjugated donkey anti-mouse secondary antibody (The Jackson Laboratory), diluted 1:500 in PBSTG for 2 h at room temperature. Glands were further washed in PBST for 6 × 10 min prior to mounting in Vectashield without DAPI.

### Preparation of flies for eye imaging

For light stereomicroscope imaging of adult eyes, female flies were collected from crosses and were aged 2-4 days at 25°C before being anaesthetised via carbon dioxide and then transferred to 0.6 ml Eppendorf tubes (SLS) and frozen at -20°C for a minimum of 24hrs, with most flies imaged within one week after freezing. To image the eyes, flies were positioned facing left-to-right, with the central part of the eye as horizontal as possible, facing the objective lens.

### Live-cell imaging

Live-cell imaging was undertaken at room temperature and broadly followed the methodology used in Singh et al., 2025. A Leica Thunder inverted wide-field microscope (Leica) was employed, at 100x magnification (Leica HCX PL FLUOTAR NA 1.3, oil objective) with a K8 sCMOS camera. Four SCs were analysed per MAG (two per lobe) for ≥10 individual virgin males. The images acquired were typically z-stacks spanning a depth of 8–12 μm with a z-distance of 0.2 μm.

Thunder technology (Leica) with small volume computational clearing (SVCC) was applied to enhance contrast and eliminate out-of-focus blur. SC morphology was visualised using FLURO-Bright Field imaging. The LED settings were as follows: GFP, 475 nm excitation at 40% laser intensity; YFP, 510 nm excitation at 42% laser intensity; and RFP, 550nm excitation at 30% laser intensity. Live MCs in the central region of MAGs were imaged similarly, using the brightfield channel to identify MC nuclei in the epithelial layer.

### Fixed-cell imaging

Fixed samples were imaged on a Zeiss LSM980 with Airyscan 2 Super-resolution upright laser scanning confocal microscope equipped with 10x (Zeiss 0.45 NA; dry) and 40x (Zeiss 1.30 NA; oil; Zeiss immersion oil, refractive index 1.518) objectives.

The two lobes of the MAG were imaged for ≥10 individual virgin males. High resolution images of the MAG’s main cells and lumen were acquired from the central region of each lobe employing the 40x objective with 1x zoom on the ZEN bluesuite Software (Zeiss), using GFP, 488 nm excitation at 2% laser intensity; DAPI, 345 nm excitation at 2% laser intensity; and mCherry, 587 nm excitation at 2% laser intensity.

For MC images, the epithelial layer was identified using the DAPI nuclear marker, which was present in the Vectashield mounting medium, and one image was taken through the middle of these nuclei for each lobe for ≥10 individual virgin males. For lumen images, a z summation was performed that was centred around the middle of the MAG lumen for both MAG lobes from ≥10 individual virgin males.

For antibody-stained glands, images were captured at the distal tip of each MAG lobe, using the 10x and the 40x objectives with 1x zoom on the ZEN bluesuite Software (Zeiss)

### Eye imaging

Brightfield images used for qualitative analysis were illuminated by two external LEDs directed either side of the eye. The GFP excitation wavelength and GFP filter on the camera was used to produce images for quantitative analysis; this provided better contrast of ommatidia and boundaries, by allowing the detection of autofluorescence in the eyes. Images were captured manually in multiple z-planes using a Leica model MSV269 stereomicroscope with CHROMYX HD camera attachment, at 8X magnification. Between 10-20 consecutive images of each eye were collected, which were then stacked using Zerene Stacker (Zerene Systems, Richland, WA) to produce a composite image of each eye. This programme uses an algorithm to combine images with different focal points into one image with an extended depth of field, providing a three-dimensional view of the eye.

### Analysis and parameters

The analysis of living SCs and MCs in MAGs employed the approaches previously used by Singh et al., 2025.

### DCG phenotypes and number of mature compartments

Deconvolved images were analysed using Fiji/ImageJ. Channels were merged to create composite images and the numbers of intact DCGs marked by GFP-MFAS and DCG-containing compartments were quantified manually using the brightfield channel. Abnormal cores, which included those with mini-cores or deformed cores, were then manually counted. The mini-/deformed core phenotype was defined by the presence of multiple small cores of diameter ≥ 0.5 µm and/or a misshapen core. A mini-core phenotype was scored if the DCG was split into at least three parts of diameter ≥0.5 µm; a deformed core was recorded when the ratio of the lengths of the longest and shortest DCG axes was >1.4.

The percentage of mini-/deformed cores was calculated relative to the total number of DCGs in non-acidic compartments per SC.

To quantify compartments marked by the YFP-Rab11 fusion protein, fluorescently labelled compartments were manually examined using z-stacks for the YFP channel (Fan et al., 2020; Wells et al., 2023). The proportion of these compartments containing Rab11-positive puncta was established was used as a measure of ILV biogenesis.

### Acidification of secretory compartments

Mature (rounded) DCG compartments that are associated with acidic structures and potentially undergoing lysosomal clearance (the DCG acidification phenotype) were scored as acidified compartments. These compartments manifest different phenotypes depending on the stage of acidification. Some have a single peripheral lysosomal structure with a slightly diffuse GFP-positive DCG that has maintained its shape, while others have completely diffuse GFP-MFAS and no obvious DCG structure in brightfield. Some acidified compartments feature multiple acidic structures arranged in an arc around the compartment boundary, with or without diffuse GFP-MFAS, while others are up to 80% or more covered by acidified domains, but still retain their spherical shape. The percentage of acidified compartments per SC was determined as a proportion of the total number of both non-acidic DCG-containing compartments and acidified compartments per SC.

To calculate the lysosomal area, a freehand tool on Fiji was employed to outline and measure the area of the SC. The threshold and analyse particles tools were utilised to determine the LysoTracker Red-positive area in the complete projection of the RFP channel. The ratio of LysoTracker Red-positive area to total SC area was calculated for each SC and expressed as % lysosomal area.

### *Ex vivo* analysis of GFP-MFAS-containing main cells

For live images captured at the distal tip of the MAG, which included one SC and parts of surrounding MCs, the Rectangle tool (700 x 700 pixels) on Fiji was used to determine the area of the field of view, and the freehand tool was employed to outline and measure the area of the SC. Subsequently, the threshold and analyse particles tools were applied to measure the area covered by GFP-MFAS in the surrounding MCs using the complete projection of the GFP channel. The percentage MC area containing GFP-MFAS was calculated within the field of view, excluding the SC area.

### Analysis of GFP-MFAS-containing main cells in fixed tissue

Using a grid tool on ImageJ, a grid of 25 squares was superimposed over the MC image for the GFP-MFAS channel. In most cases, central squares 7,8,9 were analysed, as outlined below, and the mean calculated. In cases where the gland was not flat in these areas, squares 12,13,14,17,18, or 19 (from the central block of nine squares) were selected and analysed instead. This was repeated for both lobes of each MAG, and a mean calculated, with 10 individual glands scored for each experimental condition. Each square was 100 x 100 pixels. Analysis was performed using a particle tool, where the squares were thresholded with pixel threshold value set to ≥ 25 (255 maximum) at a consistent exposure, and then the following values were calculated inside each square: number of GFP-MFAS puncta, mean size of each punctum, total area of GFP fluorescence, % area covered by GFP fluorescence, mean intensity of GFP-positive pixels, integrated density (% area of fluorescence x mean fluorescence signal intensity). We also calculated % GFP-positive area/number of puncta to determine % GFP-positive area per punctum, a measure of the size of each structure containing abnormally aggregated material.

### Analysis of GFP-MFAS in MAG lumen using fixed tissue

Using the rectangle tool on ImageJ, one square (200 x 200 pixels) was marked in the central luminal region of the middle of each lobe, and the mean GFP-MFAS intensity per lobe was calculated using ImageJ mean intensity function. This was then repeated for the other lobe. The mean of these two values was calculated for each of 10 male virgin flies.

### Image processing and preparation for figures

Live-cell images for this study were prepared using deconvoluted stacks collected from the Lecia Thunder microscope. In all resulting figures, images were a single-slice composite, chosen for their optimal representation of SC morphology. The fluorescence intensity that best captured the phenotypes of interest was selected and utilised for image processing across all the live-cell data. Consistent projection of slices and settings were applied to generate images for analysis for each genotype shown in the figures. All images were cropped to identical dimensions and focused primarily on SC details.

For fixed-gland images of MCs, a single z-plane that passed through the centre of MC nuclei was acquired using the LSM980 confocal microscope. To ensure consistency in fluorescent intensity quantification, the GFP gain settings were kept constant across all genotypes

### Quantification and statistical analysis

For comparing multiple experimental genotypes with the control, we applied the non-parametric Kruskal–Wallis test followed by Dunn’s multiple comparisons post hoc test. When comparing two groups, a non-parametric Mann–Whitney test was used. These statistical analyses were performed on GraphPad Prism. All graphs displayed in the figures show the mean value for each genotype and include error bars representing the standard deviation (SD).

### *Drosophila* eye analysis

#### Image processing

Following stacking using Zerene Stacker, composite images of each eye were processed using FIJI, with an automated set-up employing various macros (see Appendix 1 for macros). The entire process is illustrated in Figure S5. An elliptical area of 576 x 624 pixels was selected at the centre of the eye. The trainable Weka Segmentation ImageJ plugin (Arganda-Carreras et al., 2017) was trained on a series of reference images of the fly eye to detect individual ommatidia and define boundaries between them, in ‘normal’ and disordered eyes. The resulting segmented probability map, converted to 8-bit, was made binary and the watershed transformation applied, to allow identification of individual ommatidia. After processing, the inbuilt FIJI Analyse Particles function generated XY coordinates of ommatidia, which were exported to Excel.

### Data analysis

The data analysis technique used to calculate regularity is depicted in Figure S5. For each ommateum in the selected area, the angle to the nearest six ommatidia was calculated. These angles were summed for each eye of a specific genotype (approximately 250 ommatidia per eye). For the reference curve, the angles for approximately 15 females of the control genotype were summed after standardising the curve by setting the value of the most commonly occurring angle to zero for each eye. The curves for individual eyes of all other genotypes were then compared to this reference curve, again after setting the value of the most commonly occurring angle to zero, and the Pearson Correlation coefficient calculated. The clear differences changed the Pearson correlation coefficient relative to the reference curve, which was then normalised using the Fisher transformation. The Fisher transformation was applied to Pearson correlations for each eye, producing normally distributed data for statistical analysis. The transformed Z-values were exported to GraphPad Prism, where comparisons were made using one-way Brown-Forsythe and Welch ANOVA tests, followed by unpaired t-tests with Welch’s correction for multiple comparisons.

## Acknowledgements

We are grateful to all the staff at the Micron Bioimaging Facility for their support. We thank Suzanne Eaton, Stephen Goodwin, Elodie Prince, Francois Karch and Linda Partridge, as well as the Bloomington and Vienna *Drosophila* Stock Centres for *Drosophila* stocks. We acknowledge the support of the BBSRC (BB/R004862/1, BB/W00707X/1, BB/W015455/1) and Cancer Research UK (C19591/A19076). For the purpose of Open Access, the author has applied a CC BY public copyright licence to any Author Accepted Manuscript (AAM) version arising from this submission.

## Supplementary Figures and Legends

**Figure S1.**
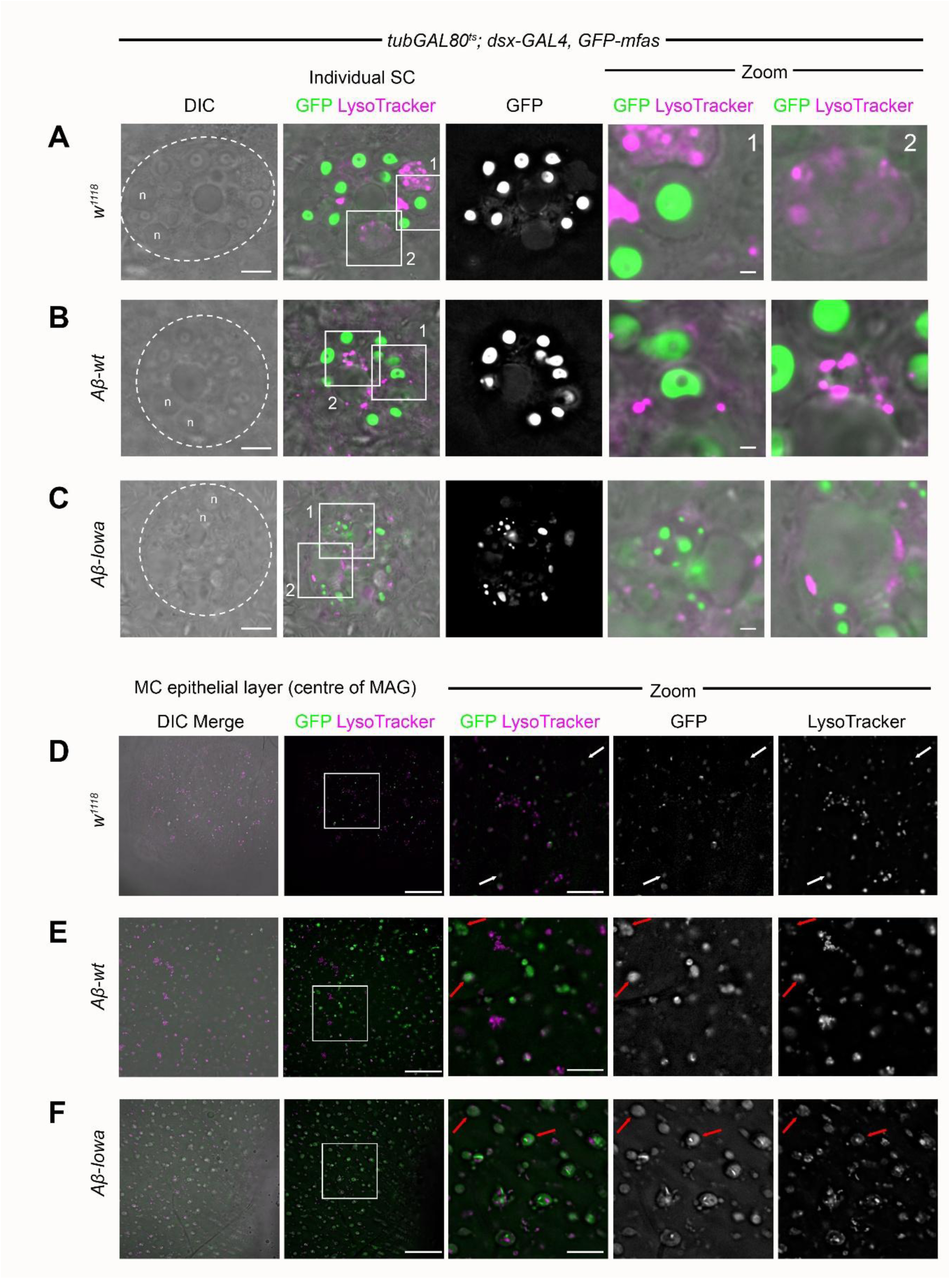
Aβ expression in SCs induces defects in MC endolysosomes (related to Figure 1) *Ex vivo* wide-field fluorescence micrographs of MC epithelial layer in middle of MAGs (see Figure 1A) from flies carrying a *GFP-mfas* gene trap and stained with LysoTracker Red. SCs either express no UAS-regulated transgene (**D**), or Aβ-wt (**E**) or Aβ-Iowa (**F**). Note when SCs express either form of Aβ, GFP-MFAS is more diffusely distributed in MC compartments and typically colocalises with larger LysoTracker Red-positive endolysosomal structures (red arrows in **E**, **F** versus white arrows in **D**). Also, Aβ-Iowa can induce formation of highly concentrated aggregated rods in some MC acidic structures. White squares mark positions of Zoom areas. Scale bars: 30 µm and 10 µm for higher magnification (Zoom) views.

**Figure S2.**
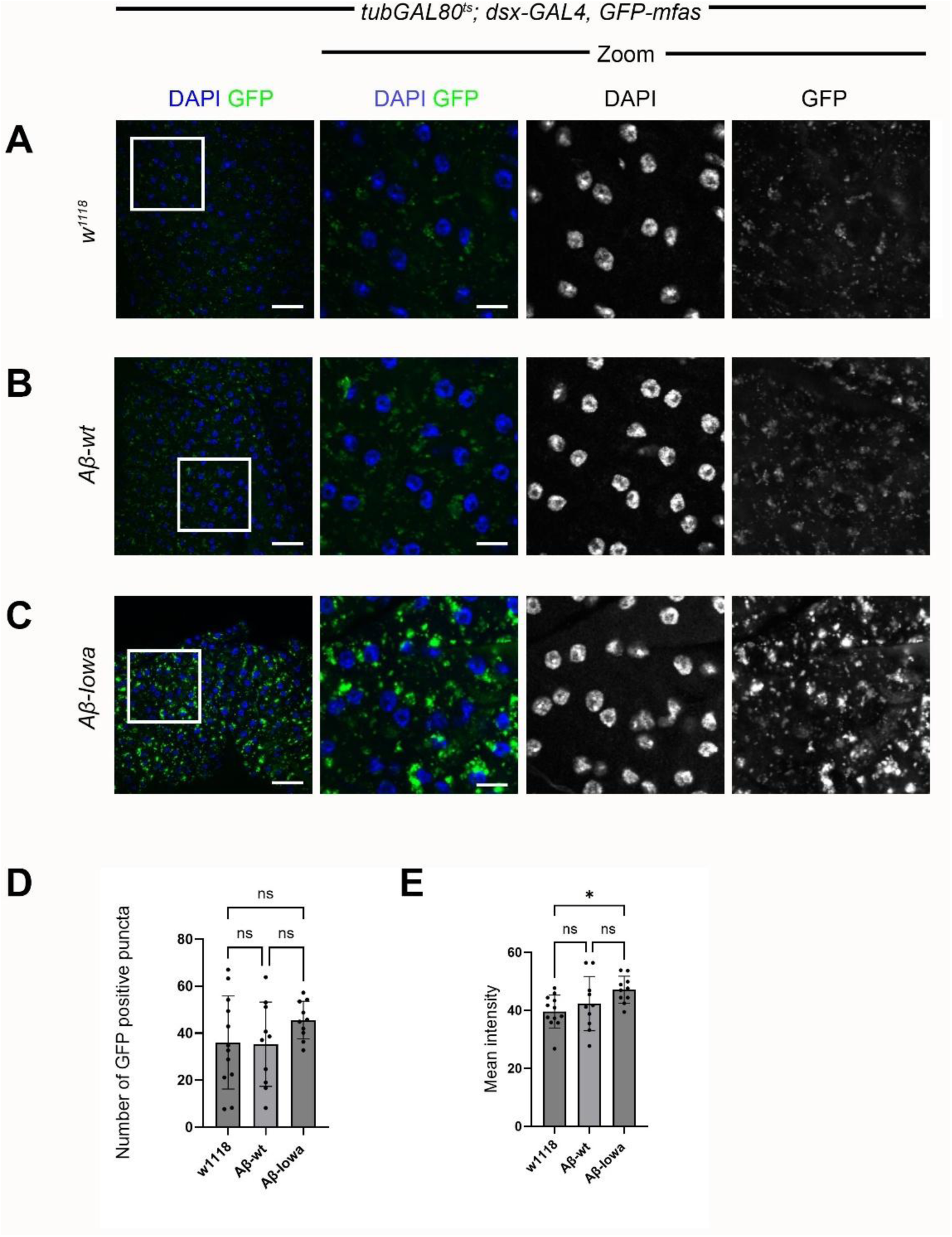
SC-expressed human Aβ isoforms induce endolysosomal defects in epithelial cells of the MAG (related to Figure 1) **A-C.** Confocal images of transverse sections through the MC epithelial layer in the proximal part of the DAPI-stained MAG from 6-day-old males expressing the *GFP-mfas* gene trap and either no other transgene (**A**), or UAS-regulated Aβ-wt (**B**) or Aβ-Iowa (**C**) specifically in SCs. **D, E.** Bar charts showing number of fluorescent puncta in field of view (**D**) and mean intensity of fluorescent puncta (**E**) in the MC epithelial layer at the centre of GFP-MFAS-expressing MAGs of the same genotypes (see also Figure 1). White squares mark positions of Zoom areas. Scale bars: 30 µm and 10 µm for higher magnification (Zoom) views.

**Figure S3.**
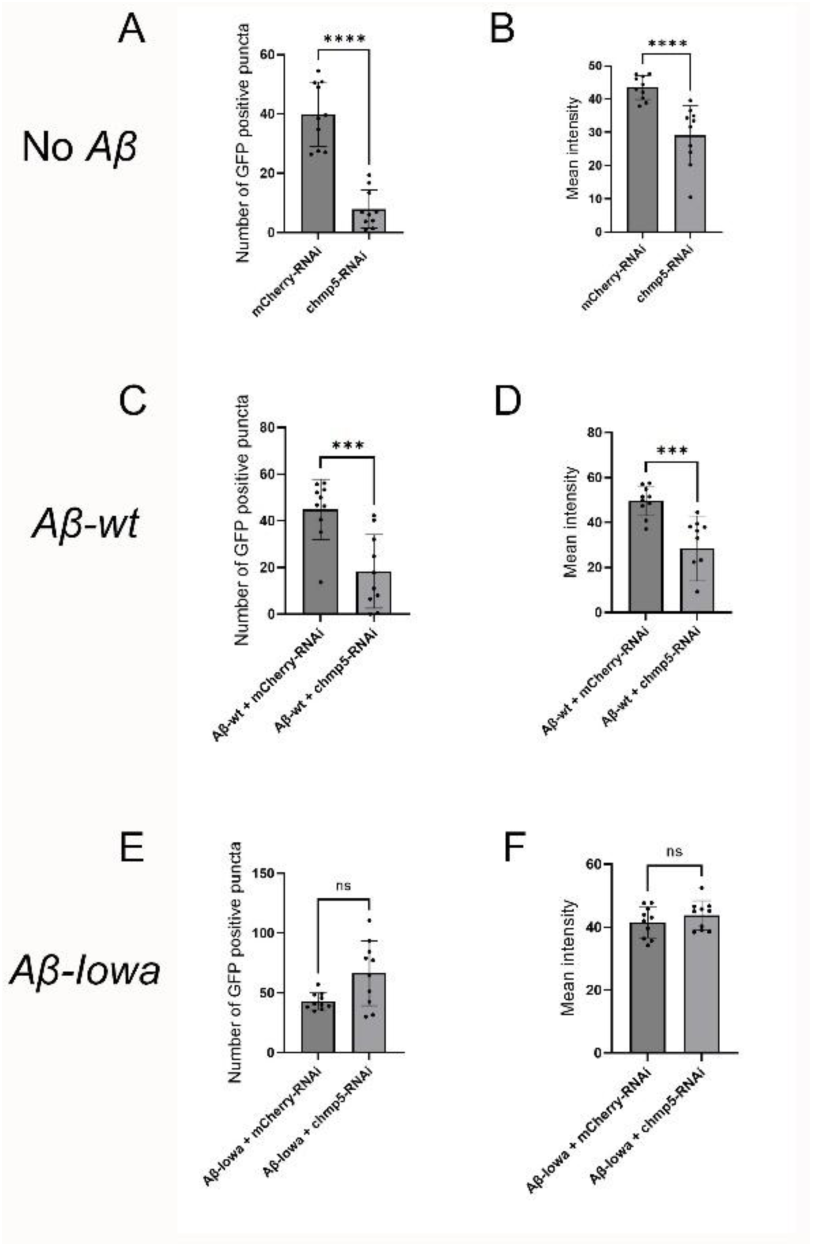
SC-specific *Chmp5* knockdown can induce multiple changes in GFP-MFAS-containing endolysosomal compartments in MCs (related to Figure 5) **A-F.** Bar charts quantifying *Chmp5* knockdown-induced changes in GFP-MFAS transfer to MCs in genotypes where SCs express no Aβ (**A**, **B**), Aβ-wt (**C**, **D**) or Aβ-Iowa (**E**, **F**) (see also data in Figure 5). There is a significant reduction in the number of fluorescent puncta in the field of view in the MC epithelial layer at the centre of the GFP-MFAS-expressing MAGs (**A**, **C**, **E**) and in mean intensity of fluorescent puncta in MCs (**B**, **D**, **F**) for Aβ-wt-expressing MAGs and non Aβ-expressing MAGs following SC-specific *Chmp5* knockdown, but not for Aβ-Iowa-expressing MAGs (**E**, **F**).

**Figure S4.**
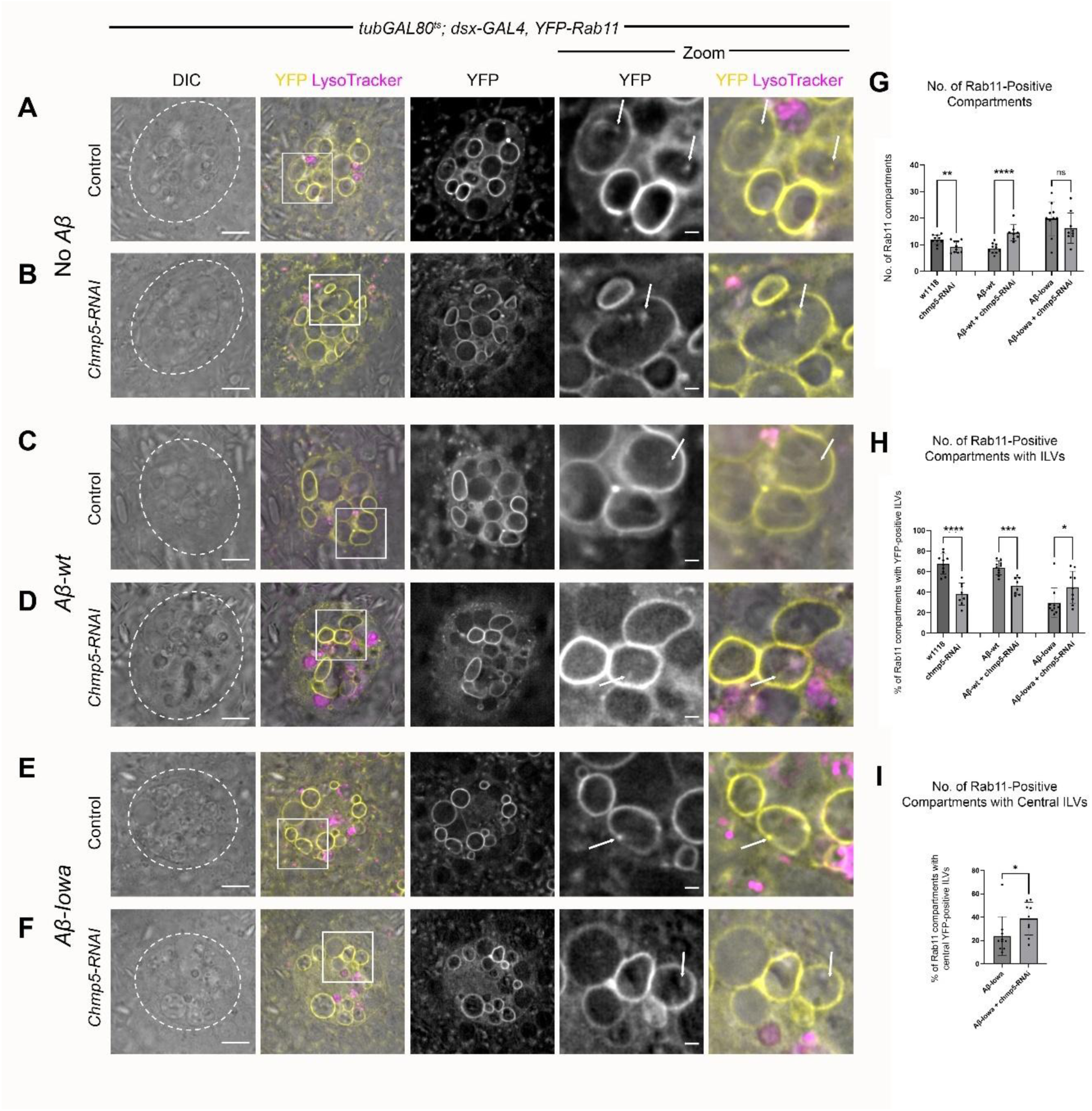
SC-specific *Chmp5* knockdown does not alter DCG compartment identity in Aβ-expressing cells (related to Figure 6) **A-F**. *Ex vivo* wide-field fluorescence micrographs of single SC (marked by dashed circle) and surrounding MCs from MAGs of flies carrying a *YFP-Rab11* fusion gene at the endogenous *Rab11* locus and stained with LysoTracker Red. SCs either express no UAS-regulated transgene (**A**, **B**), or Aβ-wt (**C**, **D**) or Aβ-Iowa (**E**, **F**) in the presence (**B**, **D**, **F**) or absence (**A**, **C**, **E**) of SC-specific *Chmp5* knockdown. Bright-field view reveals outlines of large secretory and endolysosomal compartments, as well as two nuclei (n) of these bi-nucleate cells. Zoom images focus on individual DCG compartments and highlight intra-compartmental YFP-Rab11 (arrows), which marks a subset of ILVs; *Chmp5* knockdown generally reduces the proportion of DCG compartments that contain these puncta. **G,H.** Bar charts comparing SC Rab11-positive secretory compartment number (**G**) and % Rab11-compartments containing Rab11-positive ILVs (**H**) for the genotypes shown in **A-F**. Note that the association of ILVs with the limiting membrane in SCs expressing Aβ-Iowa made it difficult to accurately assess the % of Rab11 compartments containing ILVs. *Chmp5* knockdown increased the numbers of central ILVs in Aβ-Iowa-expressing cells (**I**), so that the numbers of ILV-containing compartments appeared to increase following knockdown (**H**). White squares mark positions of Zoom areas. Scale bars: 10 µm; 2 µm in Zoom.

**Figure S5.**
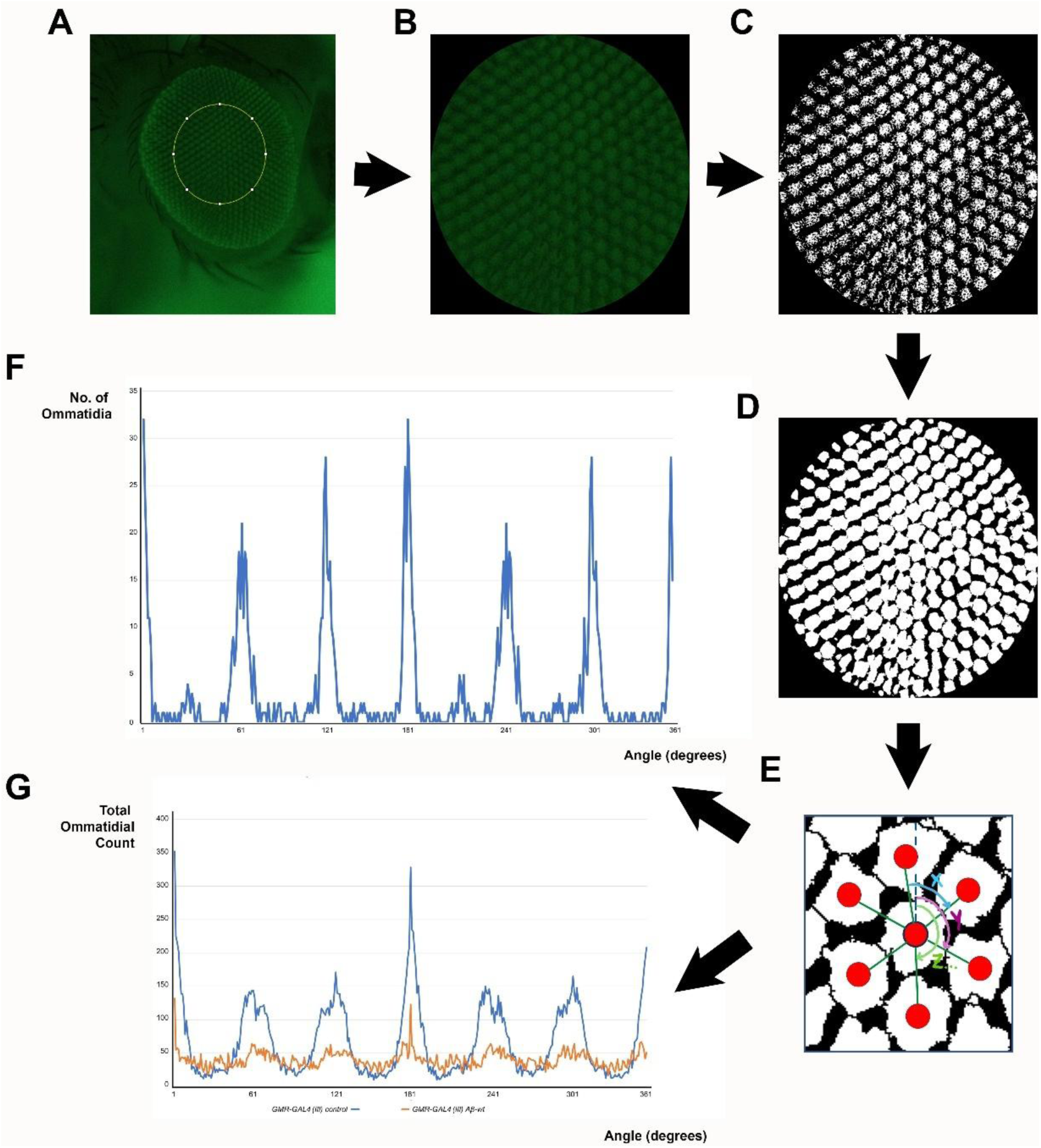
Computational analysis of ommatidial regularity (related to Figure 7) Figure illustrates analytical pipeline used to assess ommatidial regularity in the eye. **A.** One eye from each fly to be analysed is imaged using the excitation wavelength for GFP and green filter for detection of autofluorescence. An elliptical selection area (576 x 624 pixels) is placed at the centre of the eye and isolated (**B**). **C.** This selection area is converted to 8-bit greyscale and automatically thresholded in Fiji. **D.** The Trainable Weka Segmentation plugin is used to detect individual ommatidia. The FIJI built-in analyse particles function is then used to record XY coordinates for each ommatidium, which are exported to Excel. **E.** Using these data, the angles of the six closest neighbours to each ommatidium are calculated. **F, G.** These angles are summed for each eye of a specific genotype (approximately 250 per eye) and can either be summed for the control eyes used to produce the reference curve (**F**), or plotted for each eye (**G**), in both cases, after standardising the curve by setting the value of the most commonly occurring angle to zero for each eye. The graph in **G** shows data for two eyes from females, one from the *GMR-GAL4 (III)* cross to *w^1118^* control flies (blue line), the other for a *UAS-Aβ-wt; GMR-GAL4 (III)* female (orange line). The clear differences change the Pearson correlation coefficient relative to the reference curve, which is then normalised using the Fisher transformation.

**Figure S6.**
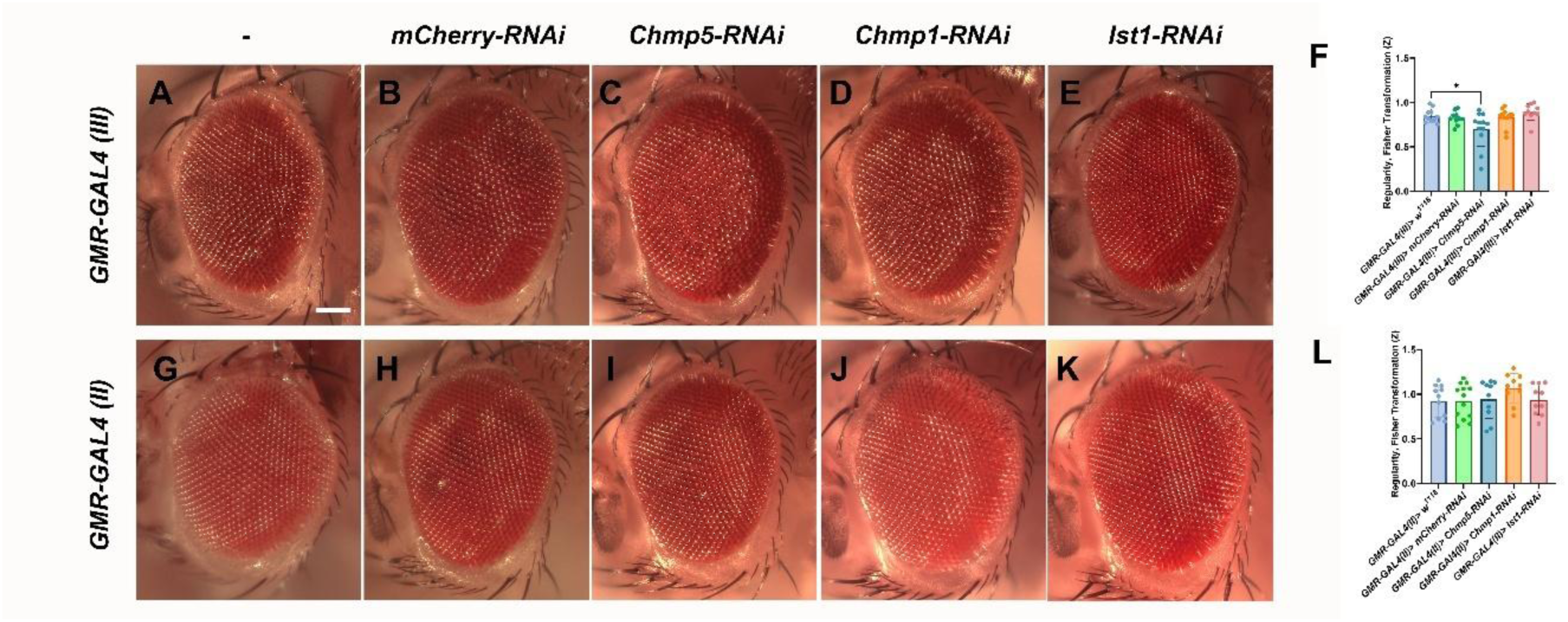
Knockdown of *accessory ESCRT-III* genes in the eye generally has no effect on ommatidial regularity (related to Figures 8 and 9) **A-E.** Stereomicroscopic images of eyes from 2-4-day-old females carrying the third chromosome *GMR-GAL4 (III)* insertion and expressing either no other transgene (**A**), *UAS-mCherry-RNAi* (**B**) or *UAS-Chmp5-RNAi* (**C**) or *UAS-Chmp1-RNAi* (**D**), or *UAS-Ist1-RNAi* (**E**). **F.** Bar chart plotting regularity score (as a Fisher Z transformation) for eyes of these genotypes relative to control *GMR-GAL4 (III)* reference curve. Note that *Chmp5* knockdown appears to produce some eyes with subtly increased disorganisation. **G-K.** Stereomicroscopic images of eyes from 2-4-day-old females carrying the second chromosome *GMR-GAL4 (II)* insertion and expressing either no other transgene (**G**), *UAS-mCherry-RNAi* (**H**) or *UAS-Chmp5-RNAi* (**I**) or *UAS-Chmp1-RNAi* (**J**), or *UAS-Ist1-RNAi* (**K**). **L.** Bar chart plotting regularity score (as a Fisher Z transformation) for eyes of these genotypes relative to control *GMR-GAL4 (II)* reference curve. Scale bar: 100 µm for all images.

## Appendix 1 Macros for processing eye images

### Automatic image crop and threshold macro

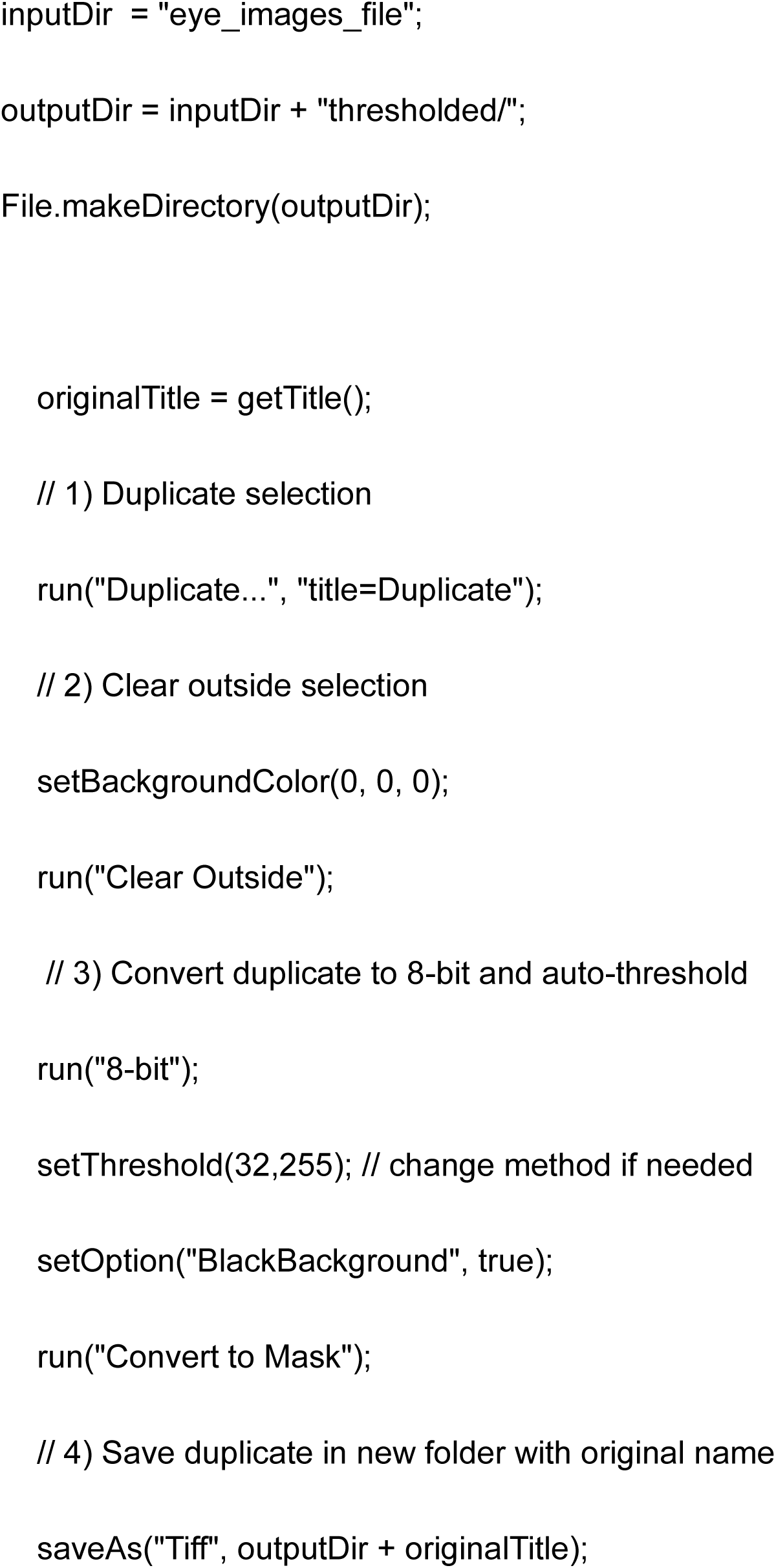
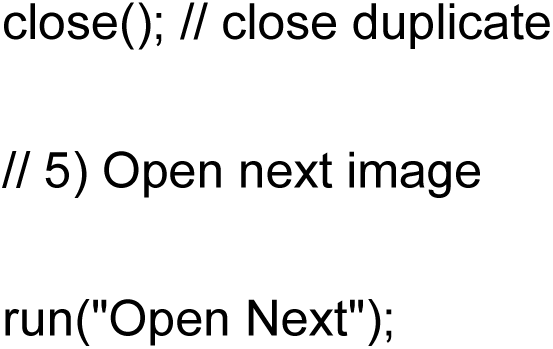

### Trainable Weka Segmentation and watershed-transformed binary image generation

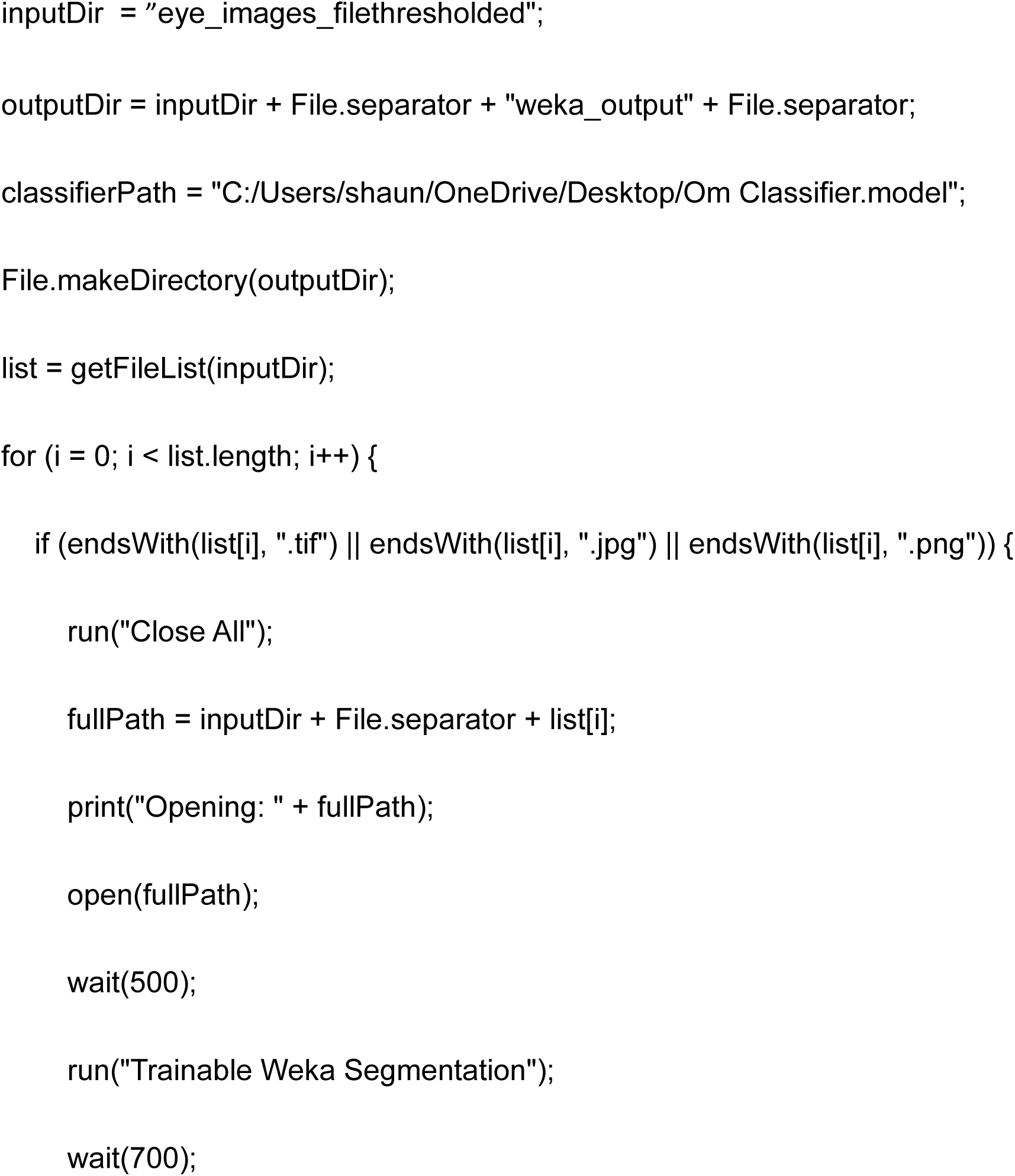
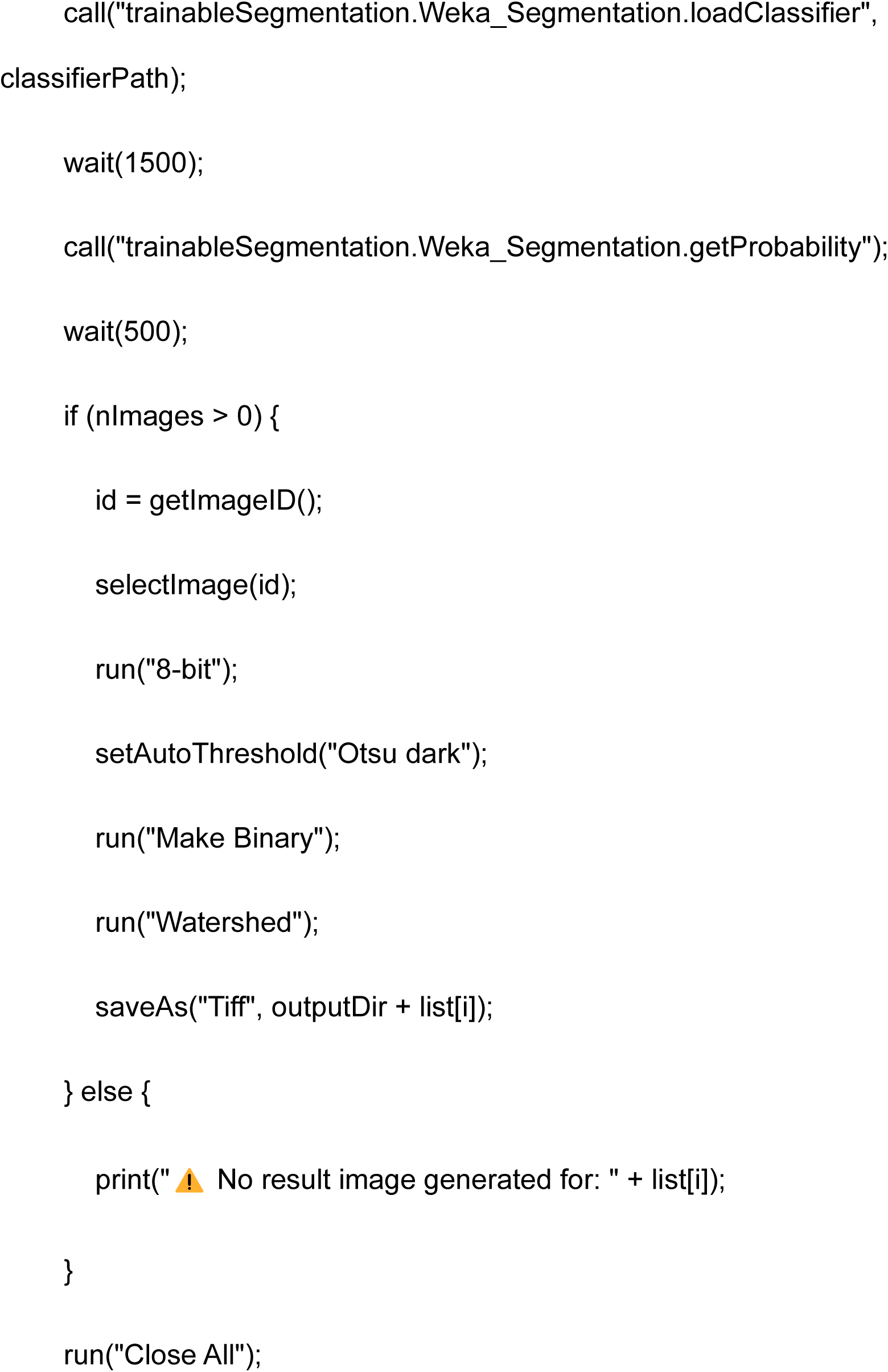

### Analyse Particles and Export to Excel

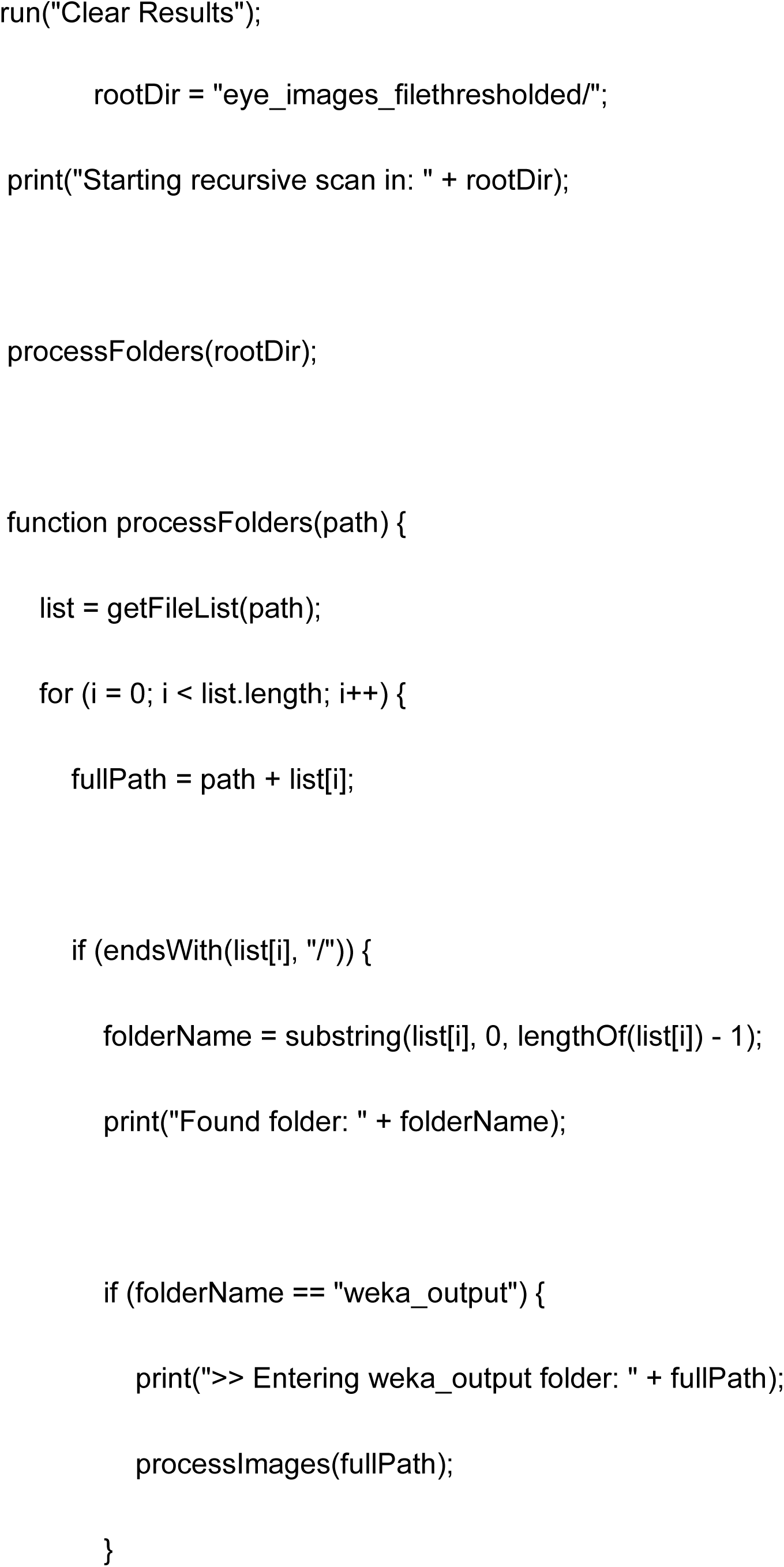
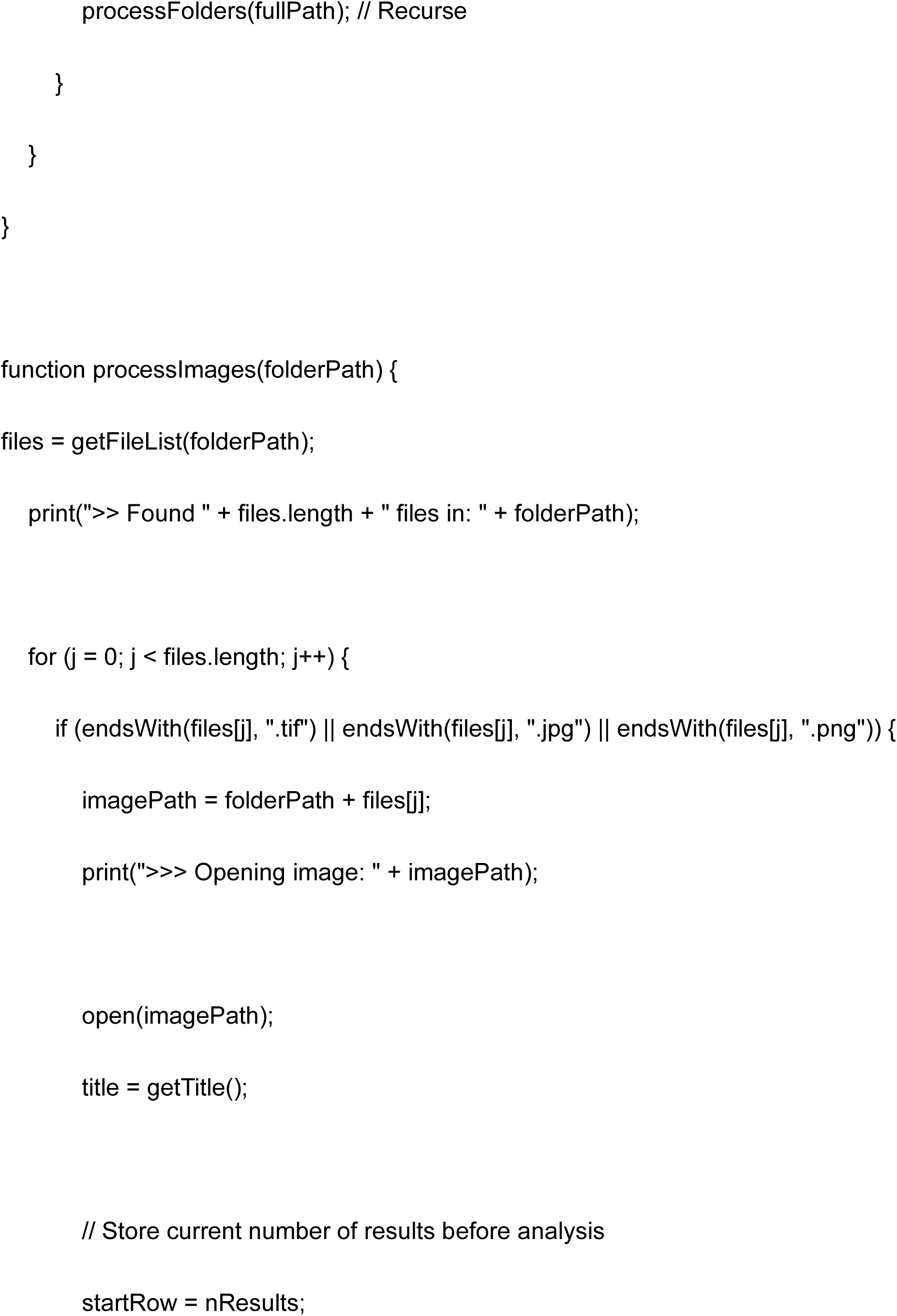
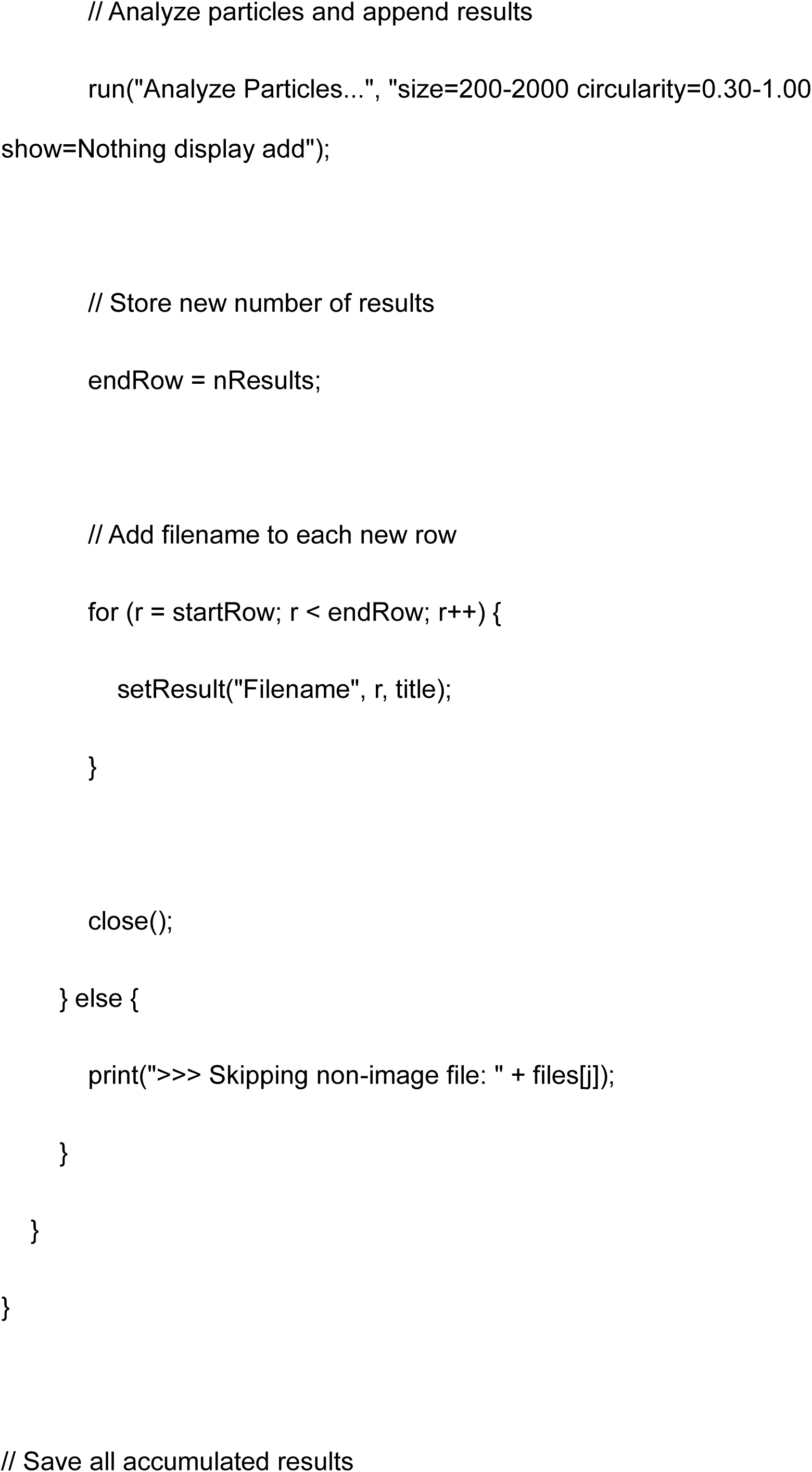
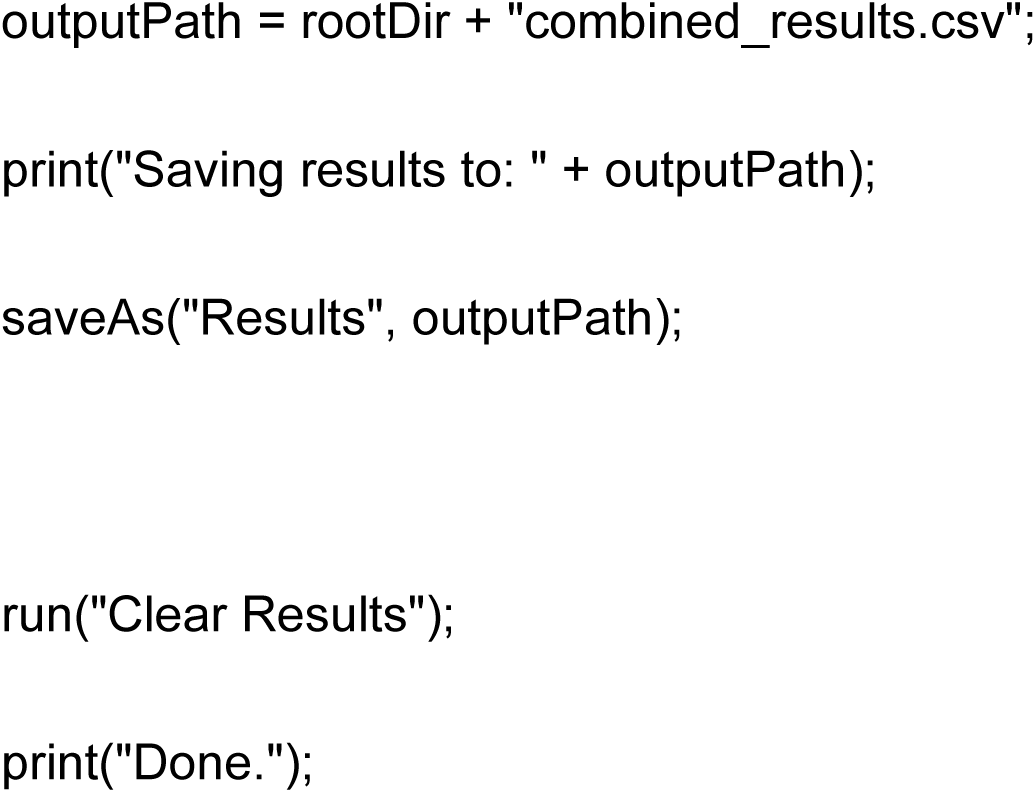

